# Molecular programming *in utero* modulates hepatic lipid metabolism and adult metabolic risk in obese mother offspring in a sex-specific manner

**DOI:** 10.1101/2021.05.26.445738

**Authors:** Christina Savva, Luisa A. Helguero, Marcela González-Granillo, Tânia Melo, Daniela Couto, Bo Angelin, Maria Rosário Domingues, Xidan Li, Claudia Kutter, Marion Korach-André

**Author notes:** Corresponding author: Marion Korach-André, Department of Medicine, Metabolism Unit, Karolinska Institute, S-141 57 Huddinge, Sweden.

## Abstract

Male and female offspring of obese mothers are known to differ significantly in their metabolic adaptation and later development of complications. We investigated the sex-dependent responses in obese offspring of mice with maternal obesity, focusing on changes in liver glucose and lipid metabolism. Maternal obesity prior to and during gestation led to hepatic insulin resistance and inflammation in male offspring, while female offspring were protected. These sex differences were explained by more efficient transcriptional and posttranscriptional reprogramming of metabolic pathways to prevent the damaging effects of maternal obesity in females compared to males. These differences were sustained later in life, resulting in a better metabolic balance in female offspring. In conclusion, sex and maternal obesity drive transcriptional and posttranscriptional regulation of major metabolic processes in offspring liver differently, explaining the sexual dimorphism in obesity-associated metabolic risk.

## Introduction

The alarming increased prevalence of overweight and obese women in reproductive age has urged the need to investigate the impact on fetal health and effects that may become evident later in life. Recent studies have demonstrated strong responses of the offspring to external factors, including nutritional, environmental and hormonal changes during the prenatal and postnatal periods^1^. Both in human and animal models, embryos exposed to overnutrition during gestation and lactation show metabolic alterations later in life, including increased risk of obesity^2,3^, impaired insulin sensitivity and glucose tolerance^4^, changes in microbiome composition^5^ and increased risk of developing fatty liver disease and hepatocellular cancer^6,7^. Therefore, understanding how maternal obesity (MO) influences offspring health is of great importance for our ability to better anticipate public health needs, and to develop practices regarding the implementation of dietary and lifestyle interventions.

Important biological and physiological differences have been observed between females and males. These differences are manifested through the sex-biased incidence of many common health problems, including cardiovascular^8^, liver^9,10^, endocrine and immune diseases^11^. Recent studies have demonstrated that female and male sex hormones, as well as sex chromosomes, contribute to the development of obesity and insulin resistance^12, 13^. Moreover, the development of age-associated diseases mostly occurs in a sex-specific manner, partly correlated with changes in sex hormone levels^13^.

Our recent studies demonstrated that even when offspring received a control diet after weaning, MO altered the hepatic and adipose lipidome of the offspring in a sex-specific manner, which may contribute to the sexual dimorphism in the metabolic adaptation later in life^14,15^. Furthermore, sex-specific responses to high calorie-diets have also been described^16^, implying that sex hormones might play a major role, although the underlying mechanisms are not well understood. Using *ob/ob* mice, we could previously show that there are sex-specific pathways of lipid synthesis in the liver which determine the molecular lipid composition, and hereby may play a key role in the sexual dimorphism of obesity-associated metabolic risk^17^. We also found that estrogen could rescue some of these affected pathways in males by controlling key genes of the lipid synthesis pathways through interaction with the nuclear estrogen receptors alfa and beta^17^.

However, the mechanisms by which MO might differently program transcriptional and posttranscriptional activities in female and male offspring have not been assessed. Therefore, we have now explored how MO affects adiposity, metabolic adaptation and hepatic lipid composition in obese female and male offspring. First, we examined the sex-specific metabolic adaptation to high fat diet in offspring, and second, whether MO affected the hepatic lipidome and transcriptome differently in female and male offspring from weaning to 6-months of age. We further evaluated if the maternal and offspring high-fat diet may determine adiposity and liver steatosis in the same individual at different time-point in life (3-months and 6-months) using magnetic resonance imaging and localized spectroscopy. We discovered that the metabolic response to MO is sex-dependent due to sex-specific transcriptional and posttranscriptional activity in the liver. MO reduced the fraction of monounsaturated and increased that of polyunsaturated lipids in male offspring, while the fraction of saturated lipids was increased at an early age in females. Finally, we also identified sex-specific hepatic lipid molecular species and transcriptional regulations associated with offspring metabolic dysfunctions in obesity.

## Results

### Maternal obesity redistributes the adipose tissue and insulin sensitivity differently in female and male offspring

Consumption of the high-fat (HF) diet by F0 dam for 6 weeks prior to mating led to a significant increase of body weight compared to the control (C) diet fed F0 dam (body weight after 6 weeks of diet: 34.6±1.8 g *versus* 22.2±0.2 g, p<0.001). F0 sires were fed the C diet throughout the study. F0 dam remained on their respective diet during pregnancy and lactation. All F1 female and male offspring received the HF diet after weaning (Fig.1a). The body weight of female offspring born from obese mothers (F-HF/HF) and those born from lean mothers (F-C/HF) was similar. In contrast, males born from obese mothers (M-HF/HF) showed significantly lower body weight than those born from lean mothers (M-C/HF) from birth until week 9 of age and thereafter gained more weight than M-C/HF, even though it did not reach significance after week 17. Males weighed significantly more than females after week 10, regardless of the maternal diet (Fig.1b; S, p<0.001). To determine if maternal obesity (MO) altered the adiposity in offspring in the short or/and long term, we defined the body fat distribution by magnetic resonance imaging (MRI) at 12-week (midterm, MID) and 25-week (endterm, END) of age in the same individual. M-C/HF had more body fat than F-C/HF at MID, which became similar in both sexes at END. M-HF/HF but not F-HF/HF accumulated less fat compared to offspring born from C diet mothers at MID but became similar at END (Fig.1c). Distribution of visceral (VAT) and subcutaneous (SAT) adipose tissue was diet- and sex-dependent. At MID, M-C/HF had more VAT than F-C/HF, but MO reduced VAT in males to the level of females (Fig.1d).

**Fig. 1.**
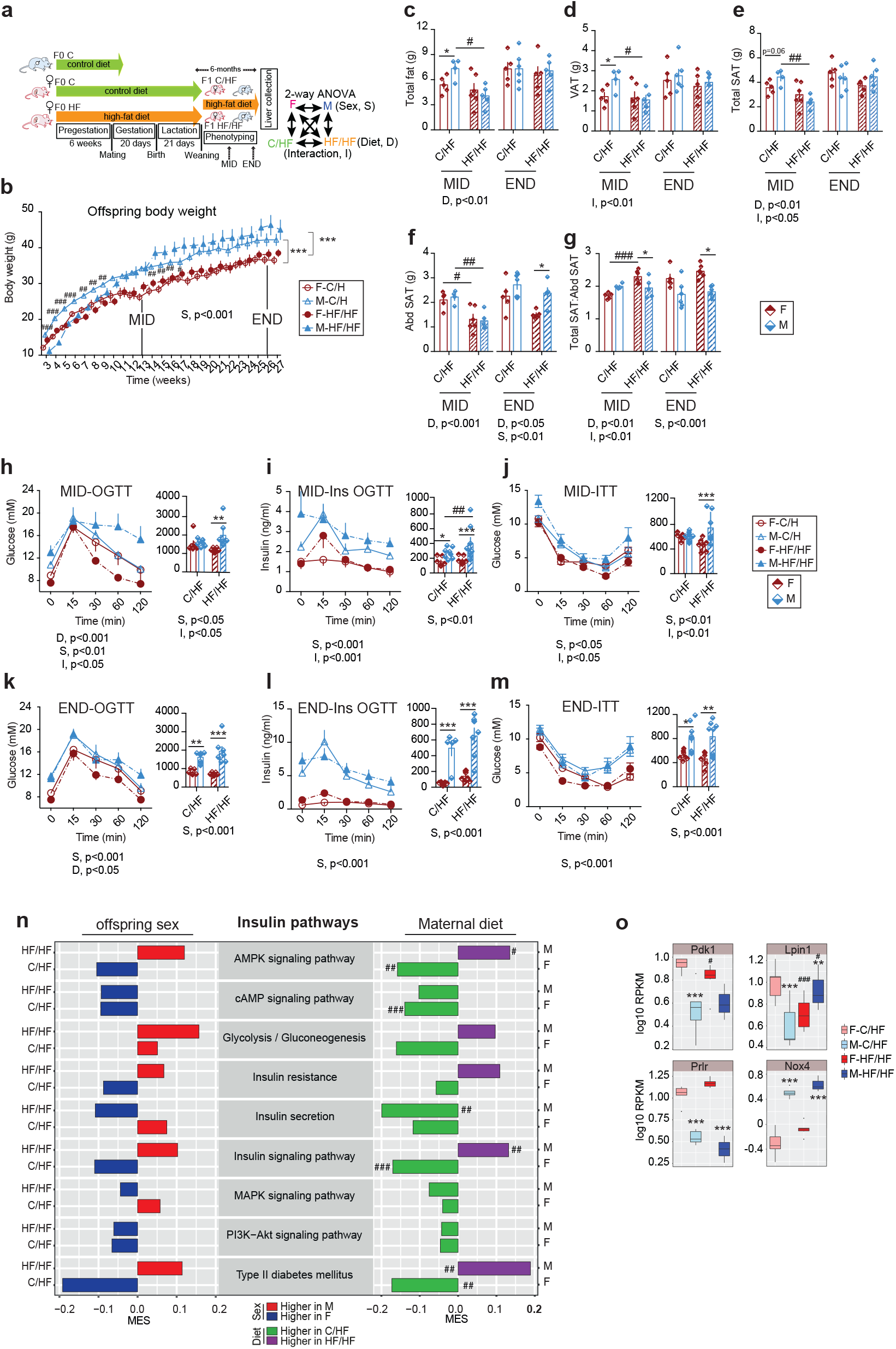
Sex-dependent physiological and transcriptional adaptations to maternal obesity in obese F1 offspring. **a** Graphic description of the study set-up. Dam-F0 were fed either the control diet (C, green arrow) or the high-fat diet (HF, orange arrow) for 6 weeks prior mating and continued on the same diet during gestation and lactation; male-F0 remained on C diet until mating. Both female and male F1 offspring remained on HF diet after weaning until sacrifice. The offspring physiological status was assessed *in vivo* at 3 months (MID) and 6 months (END) of age, using each animal as its own control. Explanatory scheme of the two-way ANOVA statistical comparisons presented on the right; **b** Time series plot of body weight in female (F, red circle; open circle in C/HF and full circle in HF/HF) and male (M, blue triangle; open triangle in C/HF and full triangle in HF/HF) offspring until sacrifice; **c-g** Bar graphs of the **c** Total fat, **d** Visceral (VAT), **e** Whole body subcutaneous (SAT) adipose tissues, **f** Abdominal SAT (Abd SAT) and **g** The ratio of total SAT on Abd SAT in F-C/HF (red open bars), M-C/HF (blue open bars), F-HF/HF (red stripped bars) and M-HF/HF (blue stripped bars) based on MRI images analysis. **h-m** Time-course of the circulating glucose levels and the corresponding insulin levels during the oral glucose tolerance test (OGTT) at, **h** and **i** MID and **k** and **l** END together with the area under the curve (AUC); Circulating glucose levels after insulin injection at **j** MID and **m** END together with the AUC; **n** Bar plot presenting the Maximum Estimate Score (MES) between sexes in C/HF and HF/HF offspring and in response to MO in F and M of the selected KEGG pathways involved in insulin and glucose metabolism; red and blue bars indicate higher expression in males and females, respectively, and green and purple bars indicate higher expression in C/HF and HF/HF, respectively; **o** Box plots of the expression level (RPKM, log_10_) of genes involved in the insulin pathways. For **b** F-C/HF (*n* =11), M-C/HF (*n* = 13), F-HF/HF (*n* = 11) and M-HF/HF (*n* = 10). For **c**–**g** F-C/HF (*n* = 7), M-C/HF (*n* = 6), F-HF/HF (*n* = 7) and M-HF/HF (*n* = 7). For **h**–**m** F-C/HF (*n* = 8), M-C/HF (*n* = 9), F-HF/HF (*n* = 7) and M-HF/HF (*n* = 6). For **n-o** F-C/HF (*n* = 5), M-C/HF (*n* = 5), F-HF/HF (*n* = 6) and M-HF/HF (*n* = 3). Data are presented as mean ± sem. Two-way ANOVA (sex (S), mother diet (D), interaction (I) between sex and diet, and (ns) for not significant) followed by Tukey’s multiple comparisons test when significant (p < 0.05). Differences between two groups (sexes, F versus M; maternal diet, C/HF versus HF/HF) were determined by unpaired t-test corrected for multiple comparisons using the Holm–Sidak method, with alpha = 5.000%. For pathway and DEG analysis we used the Benjamini-Hochberg correction with false Discovery Rate (FDR) values less than 0.1 when significant. *, M versus F and ^#^, HF/HF versus C/HF, p < 0.05; ** or ^##^, p < 0.01; *** or ^###^, p < 0.001. RPKM: Reads Per Kilobase of transcript, per Million mapped reads.

SAT is considered as the “protective” fat as it buffers extra calories intake and reduces ectopic fat accumulation^18^. We therefore investigated closer the SAT distribution in offspring. At MID, total SAT was diet- and sex-dependent but normalized at END (Fig.1e). At MID, the SAT located in the abdominal region (Abd SAT) was highly maternal diet-dependent and reduced in both sexes by MO. At END, it was higher in males than in females regardless of maternal diet (Fig.1f). The ratio between the total SAT and the Abd SAT revealed that MO redistributed SAT outside of the abdominal region in females but not in males (Fig.1g).

RNA sequencing of SAT and VAT was performed to explore if MO affected its transcriptional activity, with a special focus on browning process, inflammation and oxidative phosphorylation pathways, which play a major role in adipose tissue homeostasis^15^. Interestingly, females showed very few deregulated genes in response to MO in SAT but a highly enriched oxidative phosphorylation pathway activity, and induced *Ucp1* gene expression (marker of browning) in VAT (Suppl.Fig.S1a). Conversely, males induced significantly the expression level of a large number of genes of the inflammatory pathways, and reduced the expression level of several genes of the oxidative phosphorylation pathways in SAT (Suppl.Fig.S1a).

Changes in body fat distribution are closely correlated to metabolic disturbances. We evaluated, at the two timepoints, the glucose tolerance and insulin sensitivity in offspring by oral glucose tolerance (OGTT) and insulin tolerance (ITT) tests. At MID, glucose tolerance was highly diet- and sex-dependent, and males but not females showed impaired glucose tolerance by MO (Fig.1h). Males showed reduced insulin sensitivity (high systemic insulin levels) compared to females in both diet conditions (Fig.1i). MO impaired insulin sensitivity after insulin injection in males only (Fig.1j). At END, males showed impaired glucose tolerance together with impaired insulin sensitivity compared to females, regardless of the maternal diet (Figs.1k-m).

In sum, males but not females showed impaired insulin sensitivity with MO, possibly due to remodeling of subcutaneous fat distribution in the abdominal region associated with low-grade inflammation.

### Maternal obesity alters endocrine parameters and modulates hepatic insulin transcriptional activity in offspring

Obesity and HF diet are factors that provoke changes in circulating lipids and cytokines. While MO female offspring had unchanged total triglyceride (TG), their male counterparts displayed elevated levels (Table 1). Total cholesterol (Total Chol) levels were increased in MO offspring of both sexes, due to increased HDL-Chol (Table 1). Circulating cytokine profile was highly sex- and maternal diet-dependent. No sex differences were observed in C/HF group except for PAI-1, a strong predictor of type-2 diabetes and metabolic syndrome^19^, that was higher in males than in females in both diet groups. MO increased circulating levels of ghrelin, GIP and resistin in females to a higher level than males, all markers of improved insulin sensitivity and glucose homeostasis^20,21^. These results would indicate that MO affects positively the circulating profile in females as opposed to males that seem to stay impaired.

**Table 1.** Plasma TG, cholesterol and adipokine levels in female (F) and male (M) offspring.

| Diet | C/HF |  | HF/HF |  |
| --- | --- | --- | --- | --- |
| Sex | F | M | F | M |
| Total TG | 0.54±0.04 | 0.58±0.02 | 0.65±0.05 | 1.01±0.16<sup>#*</sup> |
| Total Chol | 2.7±0.2 | 2.4±0.3 | 3.2±0.2<sup>#</sup> | 3.6±0.3<sup>##</sup> |
| VLDL-Chol | 0.13±0.01 | 0.11±0.01 | 0.14±0.01 | 0.14±0.02 |
| LDL-Chol | 0.78±0.06 | 0.67±0.05 | 0.88±0.16 | 0.94±0.29 |
| HDL-Chol | 1.8±0.1 | 1.7±0.1 | 2.1±0.1<sup>##</sup> | 2.4±0.1<sup>###*</sup> |
| Ghrelin (mM) | 4583±341 | 5165±181 | 5074±698<sup>#</sup> | 4673±40<sup>p=0.05</sup> |
| GIP (mM) | 705±101 | 753±217 | 2716±743<sup>#</sup> | 764±29<sup>*</sup> |
| GLP-1 (mM) | 135±31 | 160±26 | 162±46 | 127±16<sup>*</sup> |
| PAI-1 (mM) | 4241±469 | 7390±997<sup>*</sup> | 4267±387 | 9633±1717<sup>*</sup> |
| Resistin (mM) | 1690±172 | 2100±380 | 2772±145<sup>#</sup> | 1742±283<sup>*</sup> |

Animals were fasted for 2h prior the blood collection. Data are presented as mean ± sem. F: female; M: male; \*, M vs F and #, C/HF vs HF/HF. \* or #, P<0.05; \*\* or ##, P<0.01; ###, P<0.001 or. For F-C/HF, n=5; for M-C/HF, n=6; for F-HF/HF, n=6; for M-HF/HF, n=4.

To explore if the observed sex-dependent metabolic adaptation to MO was associated to changes at the transcriptional level, we performed RNA sequencing in the livers of the offspring. This analysis clearly indicated that gene expression activity was sex- and maternal diet-dependent (Suppl.FigS1b). The Kyoto Encyclopedia of Genes and Genomes (KEGG) pathway analysis revealed that 32% (93/290) and 17% (48/290) of the pathways were significantly different between sexes in C/HF and HF/HF, respectively (Suppl.FigS1c; red and blue boxes). Most importantly, in about one third of the pathways (green and purple boxes) female and male offspring responded in opposite ways to MO which altered significantly 23% (68/290) of the pathways in females and 15% (45/290) in males (Suppl.FigS1c).

Since we observed impairment of insulin sensitivity in M-HF/HF, we inspected genes involved in insulin signaling pathways. There were no significant differences between the sexes (Fig.1n; left panel). However, it is interesting to note that several pathways were regulated differently between sexes between the C/HF and HF/HF groups. For example, AMPK signaling, insulin resistance and secretion, insulin signaling, and type II diabetes mellitus pathways tended to be higher expressed in females than males in C/HF but lower in HF/HF group. This suggests that MO primes insulin signaling pathways inversely in male and female offspring. Indeed, when comparing the maternal diet effect, AMPK insulin signaling, and type 2 diabetes pathways were induced upon MO in males but reduced in females (Fig.1n; right panel). In addition, MO significantly reduced cAMP signaling pathway in females and insulin secretion in males only.

In line with these results, we extracted all the differentially expressed genes (DEG) from the selected insulin pathways (Suppl.TableS1) and found four DEG known as key regulators of hepatic insulin sensitivity (*Pdk1, Lpin1, Nox4 and Prlr*) that were significantly altered by sex and by MO (Fig.1o). *Pdk1, Lpin1* and *Prlr* expressions were significantly higher in F-C/HF compared to M-C/HF while *Lpin1* was lower and *Prlr* higher in F-HF/HF compared to M-HF/HF. MO downregulated *Pdk1* and *Lpin1* expression in females but upregulated *Lpin1* in males. *Nox4* expression was higher in males than females in both diet groups (Fig.1o). Moreover, a large set of genes was differently regulated between sexes in C/HF and much less in HF/HF due to a remodeling of gene activity from the insulin pathways with MO in females only (Suppl.TableS1).

In conclusion, males showed impaired insulin sensitivity compared to females when fed a HF diet. MO impaired insulin response associated with a reduction of the signaling pathways activity at both transcriptional and post-transcriptional levels.

### Maternal obesity remodels hepatic triglyceride profile in female offspring

Obesity and insulin resistance are associated with hepatic lipid disorders, including liver steatosis, which can further develop into hepatocellular carcinoma^22^. Proton magnetic resonance spectroscopy is a prime method to track TG composition in real time (Fig.2a). Therefore, we investigated *in vivo* the TG profile in offspring livers at the two time-points (MID and END). The fraction of lipid mass (fLM) was unchanged by MO both at MID and END, with males having higher fLM than females (Fig.2b). MO induced the fraction of saturated lipids (fSL) in females at MID but not at END. At END, F-C/HF tended to have higher fSL than M-C/HF (Fig.2c). At MID, the fraction of monounsaturated lipids (fMUL) was similar in all groups, whereas MO severely reduced the fMUL in males at END (Fig.2d). At MID, MO reduced the fraction of polyunsaturated lipids (fPUL) in females, while it was unchanged in males. At END, M-HF/HF had significantly higher fPUL than F-HF/HF (Fig.2e).

**Fig. 2.**
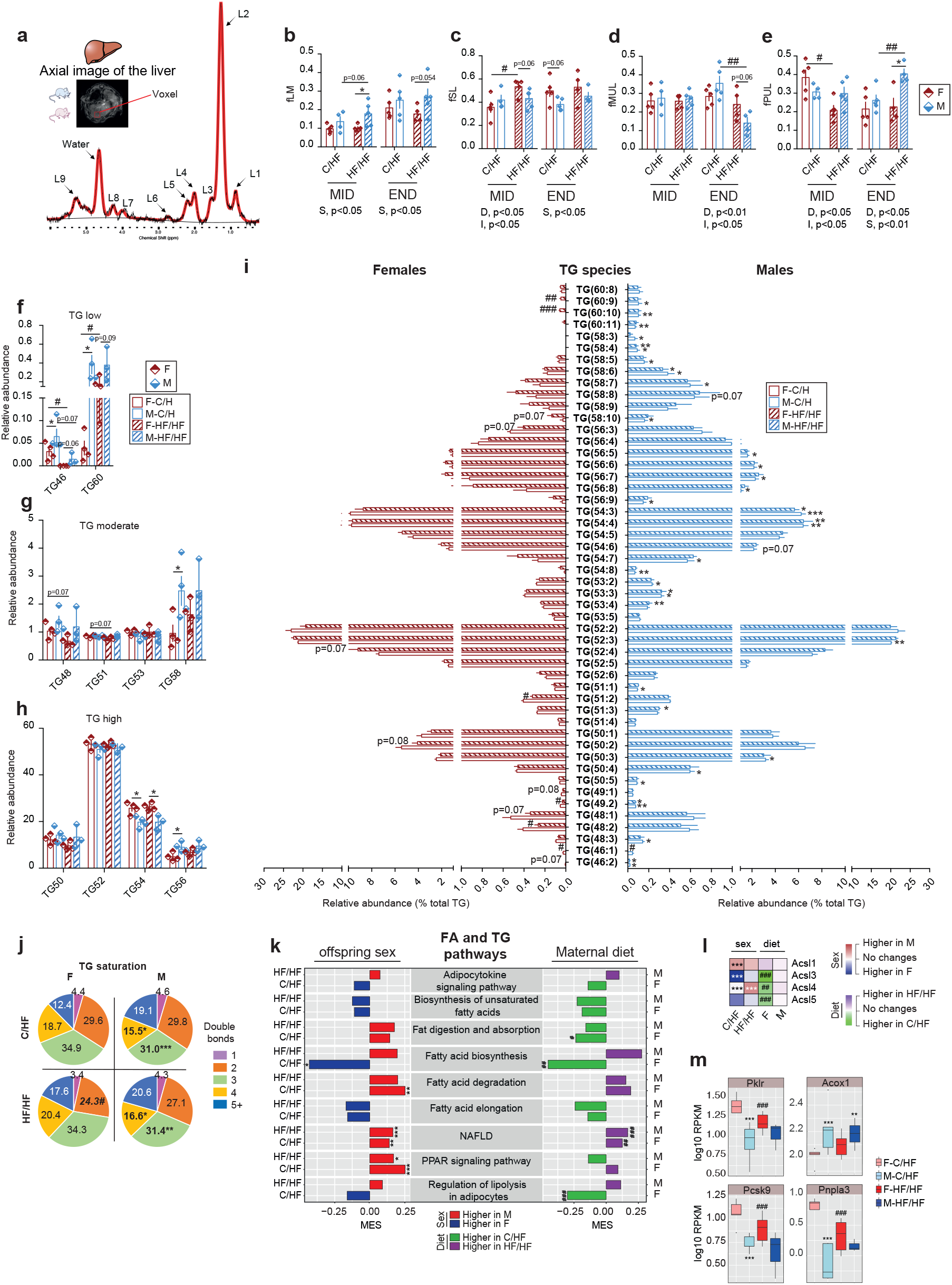
Maternal obesity adjusts the triglyceride composition in the liver of offspring and causes sex-dependent transcriptional alterations. Female (F, red bars) and male (M, blue bars) offspring born from C diet mothers (C/HF, open bars) and from HF diet mothers (HF/HF, stripped bars) at MID and END. **a** Representative axial image of the liver with single voxel spectroscopy and one representative proton spectrum used for *in vivo* quantification of the fraction of **b** lipid mass (fLM), **c** saturated lipids (fSL), **d** monounsaturated lipids (fMUL) and **e** polyunsaturated lipids (fPUL). Relative abundance of TG groups in the liver categorized as **f** low **g** moderate and **h** high abundant; **i** Bar plot of the TG species in liver extracts; **j** Pie charts showing the hepatic TG saturation profile in F and M in C/HF and HF/HF groups; **k** Bar plot presenting the MES between sexes in C/HF and HF/HF and in response to MO in F and M of the KEGG pathways involved in the FA and TG metabolism. Red and blue bars indicate higher expression in M and F, respectively and, green and purple bars indicate higher expression in C/HF and HF/HF groups, respectively. **l** Heatmap of the log2 fold change expression levels of the *Acsl* family genes and **m** Box plots showing expression (RPKM, log10) of genes involved in the FA and TG pathways. For **b**–**e** F-C/HF (*n* =5), M-C/HF (*n* = 9), F-HF/HF (*n* =5) and M-HF/HF (*n* = 5). For **f**–**j** F-C/HF (*n* =4), M-C/HF (*n* = 4), F-HF/HF (*n* =4) and M-HF/HF (*n* = 3). For **k-m** F-C/HF (*n* =5), M-C/HF (*n* = 5), F-HF/HF (*n* =6) and M-HF/HF (*n* = 3). Data are presented as mean ± sem. Two-way ANOVA (sex (S), mother diet (D), interaction (I) between sex and diet, and (ns) for not significant) followed by Tukey’s multiple comparisons test when significant (*p* < 0.05). Differences between two groups (sexes, F versus M; maternal diet C/HF versus HF/HF) were determined by unpaired t-test corrected for multiple comparisons using the Holm–Sidak method, with alpha = 5.000%. For pathway and DEG analysis we used the Benjamini-Hochberg correction with FDR<0.1, when significant. *, M versus F and ^#^, HF/HF versus C/HF, *p* < 0.05; ** or ^##^, *p* < 0.01; *** or ^###^, *p* < 0.001.

Changes in hepatic TG profile have been associated with several metabolic diseases including insulin resistance, metabolic-associated fatty-liver diseases (MAFLD) and hepatocellular carcinoma^23,24^. Therefore, we analyzed the TG composition in harvested livers using lipidomic. We found 10 TG groups that were classified as low abundant (TG low), moderate abundant (TG moderate) and high abundant (TG high). Overall, TG groups were sex-dependent with M-C/HF having more TG46, TG56, TG58 and TG60 than F-C/HF; but less of TG54 (Figs.2f-h). Interestingly, MO tended to reduce the proportion of short chain TG (TG46, TG48, TG50 and TG51) and to increase long chain TG (TG56, TG58 and TG60) in females only (Figs.2f-h). TG species comprised in each TG group were also highly sex-dependent, but MO remodeled TG species mostly in females (Fig.2i and Suppl.Figs.S2a-c). Modification of the saturation profile of hepatic TG has been correlated to several metabolic dysfunctions. Males showed lower abundance of TG-containing 3- and 4-double bonds and tended to have more TG-containing 5+-double bonds than females regardless of the maternal diet (Fig.2j). MO reduced the proportion of TG-containing 2-double bonds in females.

Fatty acids (FA) as part of TG molecules act as signaling molecules that can modulate metabolic response in obesity. FA composition was sex-dependent whereby males had higher abundance of the C20:0, C20:1ω9, C20:2ω9 and C20:3ω6 species compared to females, irrespective of the maternal diet. MO increased the abundance of C18:2ω6 and reduced C16:0 species in females (Suppl.Fig.S3a). MO increased the proportion of ω6 FA in females (Suppl.Fig.S3b). M-C/HF had globally more of the PUFA than F-C/HF; MO increased FA-containing 2- and 3-double bonds in females (Suppl.Fig.S3c). Desaturation of FA is controlled by desaturase enzymes. The desaturase activity Δ19 was unchanged between groups but Δ15 was significantly higher in females than in males in both diet conditions (Suppl.Fig.S3d). In sum, hepatic FA and TG composition is sex dependent, and MO remodeled FA and TG profiles differently in female and male offspring, which may promote sex-dependent liver dysfunctions later in life.

KEGG pathway analysis revealed that PPAR signaling and FA degradation pathways, indicative for FA breakdown, as well as NAFLD pathway were higher expressed in males than in females in both diet conditions (Fig.2k, left panel). In contrast, the FA biosynthesis pathway activity was higher in F-C/HF than in the M-C/HF but not in the HF/HF group due to a significant reduction of activity by MO in females. MO reduced fat digestion and absorption and lipolysis in adipocytes pathways activity in females while increased the NAFLD pathway in both sexes (Fig.2k, right panel). We next inspected genes involved in the selected lipid pathways and found a large number of DEG between sexes in C-HF and much less in HF/HF. These changes were explained by remodeling of gene activity by MO in females only (Suppl.TableS2). We then extracted key DEG involved in hepatic lipid homeostasis, namely the long-chain acyl-CoA synthase family members (*Acsl1/3/4/5*), the *Pklr, Acox1, Pcsk9* and *Pnpla3* genes. Their expression levels were highly sex-specific and were reduced by MO in females (Figs.2l-m). Modulation of these genes affects hepatic intracellular TG levels and the viability of these cells^25,26^.

In conclusion, hepatic transcriptional regulation of lipid pathways is sex- and maternal diet-dependent and may be a key contributor to the sexual dimorphism in obesity-associated liver disorders.

### Maternal obesity remodels hepatic phospholipid profile in offspring

In hepatic cells, phospholipids (PL) comprise the most abundant lipid class. PL are found in the plasma membrane and intracellular organelles, and the lipidome of each organelle may be remodeled by extra- and intra-cellular stimulations that may also affect lipid trafficking across the membrane and the organelles. We comprehensively profiled hepatic PL using a LC-MS lipidomic approach. Principal component analysis separated the PL classes into two distinct groups clustered by sexes (Fig.3a), which indicates that PL profile is strongly sex dependent. Four major subclasses of PL were found, phosphatidylcholine (PC), lysoPC (LPC), phosphatidylethanolamine (PE) and lysoPE (LPE) (Fig.3b and Suppl.Figs.S4a-b). The relative abundance of the total PC, PE and LPC classes was similar between sexes in both maternal diet conditions, but males had more LPE than females regardless of the maternal diet (Fig.3c and Suppl.Figs.S4a-b). However, F-C/HF had higher relative level of PC30, PC40 and PC42 classes and lower level of PC36 class than M-C/HF. MO increased the relative level of PC34 class to a higher level in males than in females, and increased the PC40 class in females to a higher level than males (Fig.3c). No differences were observed between sexes in PE classes regardless of maternal diet, but males overall tended to have higher levels of PE than females (Fig.3d). In the PC and PE classes, 15/30 PC and 8/25 PE species had different abundancies between sexes in C/HF group. MO reduced these differences considerably, to 5/30 for PC and 2/25 for PE. This occurred mainly through remodeling PC and PE species in females (Suppl.Figs.S4c-d). PC and PE saturation profiles were sex-dependent in C/HF offspring, and MO abolished the sex differences (Fig.3e). Interestingly, males tended to have higher relative levels of the LPC species, and had higher levels than females of most of the LPE species regardless of the maternal diet (Figs.3f-g and Suppl.Figs.S4e-f). The saturation profile was highly variable between sexes in C/HF but not in HF/HF group (Figs.3h-i), consonant with remodeling in female offspring by MO.

**Fig. 3.**
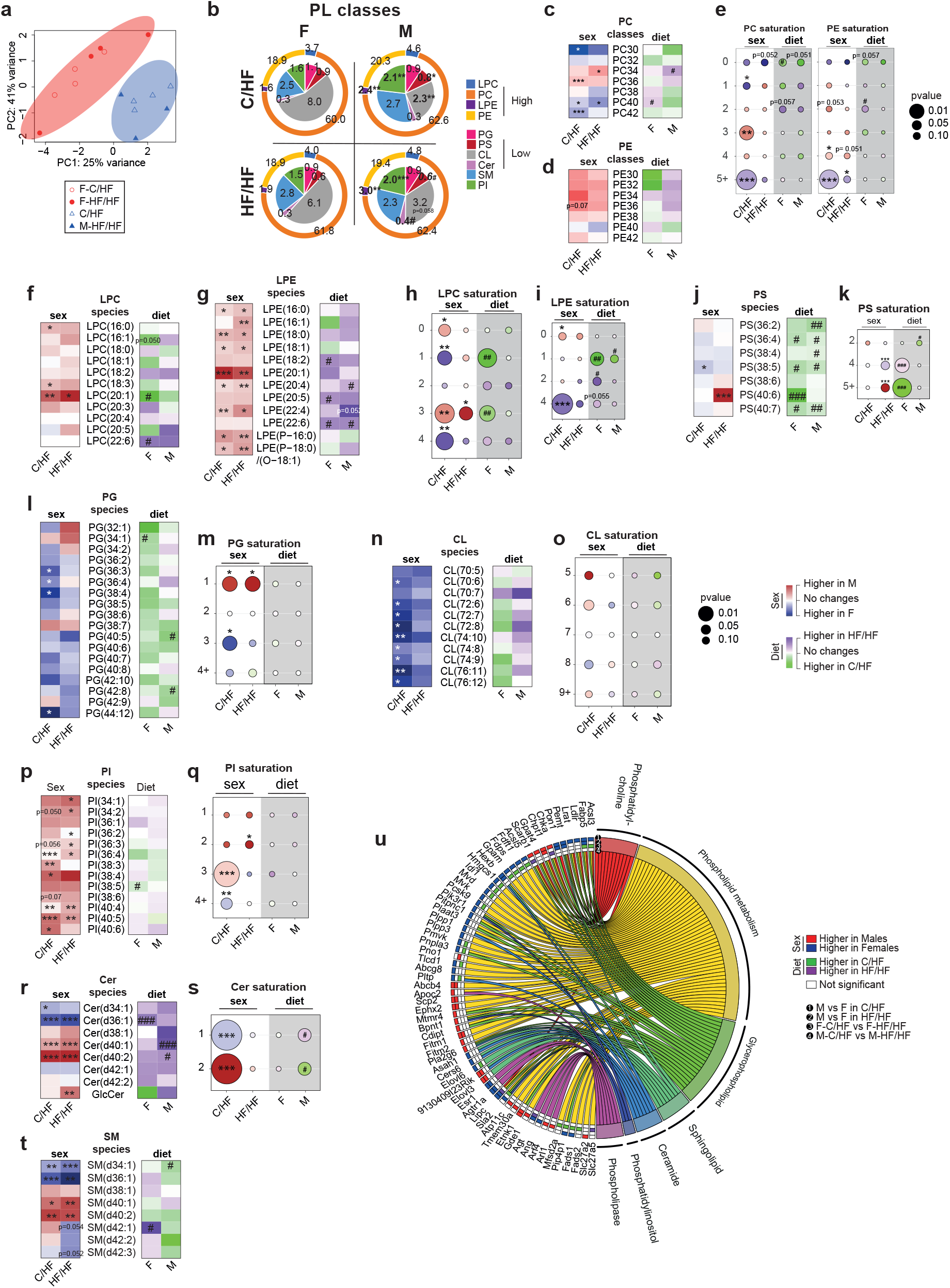
Hepatic phospholipid composition is sex-dependent in offspring regardless of maternal diet. **a** Principal component analysis (PCA) plot of the phospholipid profile in liver extracts; **b** Chart representing relative abundancies of all PL classes categorized in low (inner layer of the chart) and major classes (outer part of the chart); Heatmaps presenting the log10 fold change between sexes in C/HF and HF/HF (red and blue boxes) and in response to MO in F and M (green and purple boxes) of the **c** phosphatidylcholine (PC) and **d** phosphatidylethanolamine (PE) lipid classes; **e** Bubble charts showing the log10 fold change difference of the saturation profile in PC and PE classes between sexes (C/HF and HF/HF columns, white background) and in response to MO (F and M columns, grey background); Heatmap presenting the log10 fold change between sexes in C/HF and HF/HF and in response to MO in F and M of the **f** lysoPC (LPC), **g** lysoPE (LPE), **j** phosphatidylserine (PS), **l** phosphoglycerides (PG), **n** cardiolipin (CL), **p** phosphatidylinositol (PI), **r** ceramide (Cer) and **t** sphingomyelin (SM) lipid species; Bubble charts showing the log10 fold change difference of the saturation profile in **h** LPC, **i** LPE, **k** PS, **m** PG, **o** CL, **q** PI and **s** Cer classes between sexes (C/HF and HF/HF columns, white background) and in response to MO (F and M columns, grey background); **u** Chord graph presenting the genes associated with phospholipid pathways. The significant differential expression of each gene based on log2 fold change is presented as i) red boxes for upregulation in M, ii) blue boxes for upregulation in F, iii) green boxes for upregulation in C/HF iv) purple boxes for upregulation in HF/HF and v) white boxes when not significant. For **a-t** F-C/HF (*n* =4), M-C/HF (*n* = 4), F-HF/HF (*n* =4) and M-HF/HF (*n* = 3). For **u** F-C/HF (*n* =5), M-C/HF (*n* = 5), F-HF/HF (*n* =6) and M-HF/HF (*n* =3). For gene expression analysis (**3u**) we used the Benjamini-Hochberg correction with FDR<0.1, when significant. Data are presented as mean ± sem. Differences between two groups (sexes, F versus M; maternal diet C/HF versus HF/HF) were determined by unpaired t-test corrected for multiple comparisons using the Holm–Sidak method, with alpha = 5.000%. *, M versus F and ^#^, HF/HF versus C/HF, *p* < 0.05; ** or ^##^, *p* < 0.01; *** or ^###^, *p* < 0.001.

Other low abundant subclasses of PL were detected by LC-MS, namely phosphatidylserine (PS), phosphatidylglycerol (PG), cardiolipin (CL) and phosphatidylinositol (PI) and two sphingolipids, ceramides (Cer) and sphingomyelin (SM) (Fig.3b). The PS class, and the relative levels of 5/7 and 3/7 PS species were significantly reduced by MO in males and females, respectively (Fig.3j and Suppl.Figs.S5a-b). The PS saturation profile was affected by MO and was sex-dependent in HF/HF (Fig.3k). PG classes and species were higher in females than males in C/HF, but MO tended to reduce most of the PG species in females to the level seen in males. The PG saturation profile was sex-dependent, with no effect of maternal diet (Figs.3l-m and Suppl.Figs.5c). CL class and species were more abundant in females than males, especially in C/HF group. MO tended to reduce CL level in females with no differences in the saturation profile between all groups (Figs.3n-o and Suppl.Fig.S5d). PI classes and species were more abundant in males than in females regardless of the maternal diet, with males having more of the PI-containing 3- and 2-double bonds than females in C/HF and in HF/HF respectively. F-C/HF showed more of the PI-containing +4-double bonds than M-C/HF (Figs.3p-q and Suppl.Fig.S5e).

The Cer class was induced by MO in both sexes, with females having more of the Cer(d34) and Cer(d36) and less Cer(d40) classes than males in both maternal diet groups (Fig.3r and Suppl.Fig.S5f). Of note, the glucosylceramide (GlcCer) species were reduced and induced by MO in females and males respectively, leading to higher abundance in males. F-C/HF had more of the ceramides containing 1-double bond and less of those containing 2-double bonds than M-C/HF; no differences were observed between sexes in the MO group (Fig.3s). Females and males had similar relative SM abundance but contained different SM species in both maternal diet groups (Fig.3t and Suppl.Fig.S5g). DEG analysis of RNA-seq data established that sex is a major regulator of the PL pathways at the transcriptional level and demonstrated that MO remodeled gene expression in females only (Fig.3u).

In conclusion, PL classes and species are mainly dependent on sex, while the maternal diet in particular influences the LPE, PS and Cer classes. Overall, females have more PG and CL, and less lysoPL and sphingolipids, than males. These sex-specific classes of PL could be attributed to sex-dependent transcriptional activity of major genes involved in the PL synthesis pathways. The sexual dimorphism in the PL profile could contribute to the sex-dependent metabolic adaptation to MO by modulating transmission of biological signals across the cell and lipid droplet membranes.

### Transcriptional and posttranscriptional regulation of metabolic pathways in offspring liver is sex- and maternal diet-dependent

Hematoxylin-eosin-stained liver sections from female and male offspring showed that males had higher number and lager size lipid droplets compared to females (Fig.4a). TG and PL are central in the control of fatty liver diseases^23,27^. In addition, dysfunctional TG and PL (such as saturated PL) may initiate ER stress and inflammation^28^. To investigate the transcriptional hepatic metabolic regulation in offspring that might account for sexual dimorphism in obesity, we first performed a genome-wide differential gene expression analysis in response to MO in females and in males (Figs.4b-d), and between sexes in C/HF and HF/HF (Figs.4e-g). This revealed that females showed more DEG than males (325 *vs* 33) in response to MO and only four DEG were shared between sexes (*Cyp2c37, Cyp2c50, Cyp2c54* and *Sult2a8*). These four genes are key regulator of hepatic lipid and energy metabolism, and were all up-regulated by MO in both sexes (Suppl.Fig.S6a). Compared to females, males showed higher expression levels of *Sult2a8* and *Cyp2c54* and a lower expression level of *Cyp2c37* in both diet. About half of the DEG were up- and half were downregulated by MO in both sexes (Figs.4c-d). When we compared sex differences for each maternal diet conditions, we observed about half of the DEG between sexes (46%) exclusively in C/HF group and about one third (32%) exclusively in HF/HF group. Only 22% of the DEG were shared between C/HF and HF/HF (Fig.4e). Among the DEG about half were down- and half were upregulated between sexes (Figs.4f-g).

**Fig. 4.**
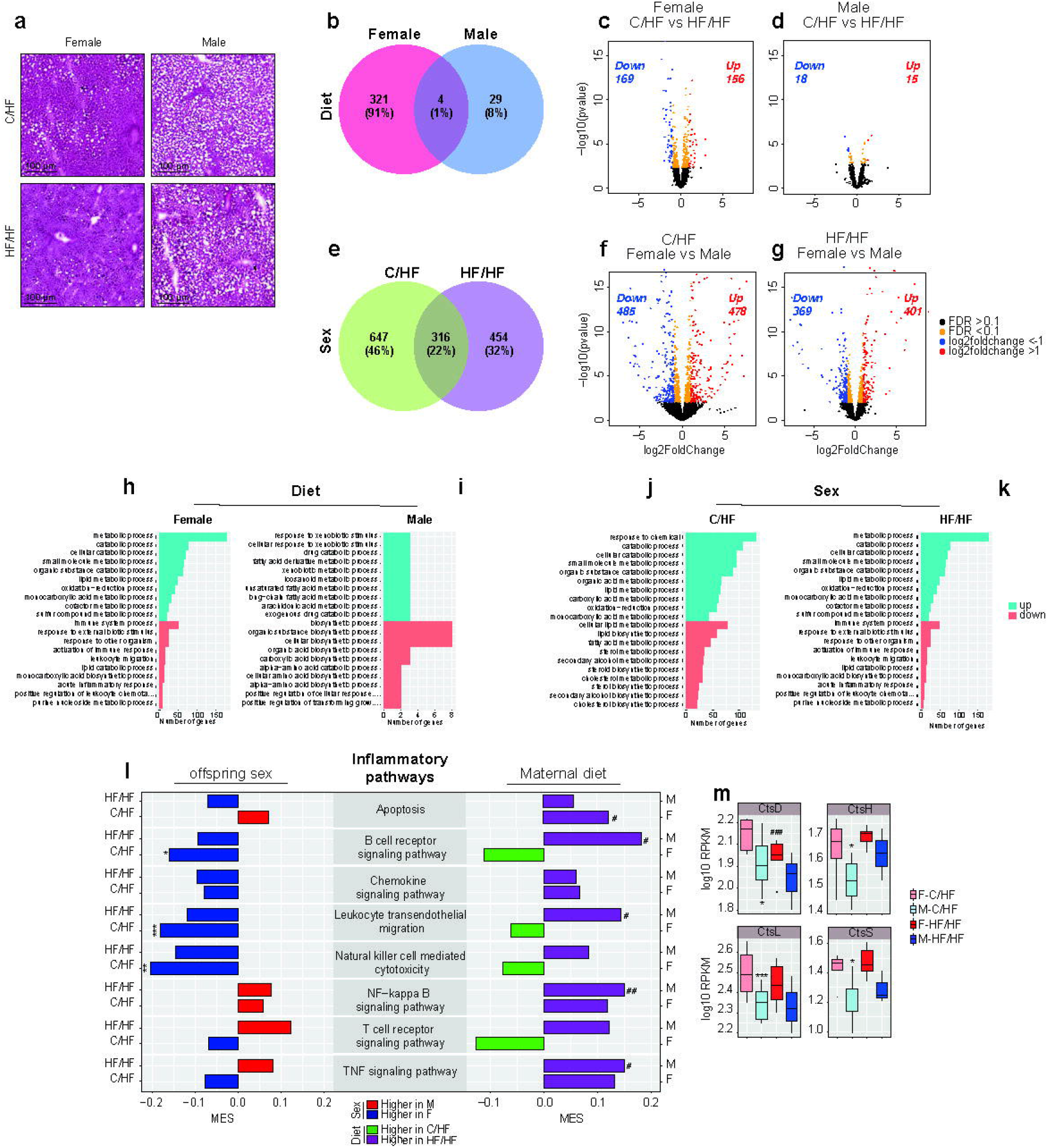
MO promotes hepatic inflammatory response in obese male’s offspring. **a** Hematoxylin-eosin (H&E) staining of frozen liver sections from C/HF and HF/HF offspring; **b** Venn diagram of the DEG in response to MO, in females (F) and males (M) and; Volcano plots of the DEG in response to MO in **c** F and **d** M; **e** Venn diagram of DEG between sexes in C/HF and HF/HF and; Volcano plots of the DEG between sexes in **f** C/HF and **g** HF/HF; Top 10 significantly up and top 10 significantly down enriched biological GO terms in response to MO in **h** F and **i** M; and between sexes in **j** C/HF and **k** HF/HF; **l** Bar plot presenting the MES between sexes in C/HF and HF/HF and in response to MO in F and M of the KEGG pathways involved in inflammation. Red and blue bars indicate higher expression in M and F, respectively and, green and purple bars indicate higher expression in C/HF and HF/HF groups, respectively. **m** Box plots showing expression (RPKM, log10) of selected genes involved in the inflammatory pathways. For volcano plots: Significantly upregulated (log2FC>1) and downregulated (Log2FC<-1) genes are presented as red and blue dots, respectively. Orange dots indicate the genes that are significantly changed (FDR<0.1). Black dots indicate not significant (FDR >0.1). For **a** F-C/HF (*n* =3), M-C/HF (*n* = 3), F-HF/HF (*n* =4) and M-HF/HF (*n* =2). For **b**–**m** F-C/HF (*n* =5), M-C/HF (*n* = 5), F-HF/HF (*n* =6) and M-HF/HF (*n* =3). Data are presented as mean ± sem. Differences between two groups (sexes, F versus M; maternal diet C/HF versus HF/HF) were determined by unpaired t-test corrected for multiple comparisons using the Holm–Sidak method, with alpha = 5.000%. For **h-k** significance was determined using Fisher’s Exact test with P values ≤ 0.05. For pathway and DEG analysis we used the Benjamini-Hochberg correction with FDR<0.1 when significant. *, M versus F and ^#^, HF/HF versus C/HF, *p* < 0.05; ** or ^##^, *p* < 0.01; *** or ^###^, *p* < 0.001.

We performed GO enrichment analysis for up- and downregulated genes and extracted the top 10 enriched GO terms for up- and downregulated genes. In females, MO upregulated metabolic and catabolic processes and downregulated immune and inflammatory processes (Fig.4h). In males, MO upregulated xenobiotic and fatty acid metabolic processes and down-regulated biosynthetic processes (Fig.4i). When comparing sexes in the C/HF group, we found increased catabolic processes and lipid metabolism pathway activity and decreased lipid and steroid biosynthetic activity in males compared to females (Fig.4j). In HF/HF, males showed increased metabolic and catabolic processes and a decrease in immune and inflammatory pathways activity compared to females (Fig.4k). Altogether these results indicate that MO alters the liver transcriptome much more in females than in males. We confirmed that MO reprograms the transcription of genes in offspring liver in a sex-dependent manner.

Given the sex-difference observed for the content of hepatic lipid droplets, we explored pathways involved in inflammation to compare the effects of sex and maternal diet (Fig.4l). This analysis unveiled that the activity of inflammatory pathways was higher in F-C/HF than in M-C/HF. These differences were abolished by MO, whereas it induced inflammatory pathway activity mainly in males. In contrast, females significantly induced gene expression program related to apoptosis pathways, and we observed a trend encompassing reduced gene expression of B cell receptor, leukocyte migration, natural killer and T cell receptor signaling pathways. These results indicate that there is a sex-dependent regulation of inflammatory pathways in livers of offspring, and that MO modulates pathways differently between sexes. We then extracted the DEG in all the selected inflammatory pathways and could show that MO altered the expression of a very few genes in female livers only (Suppl.TableS3). Among those, we found genes belonging to the cathepsin (*Cts*) family that drive liver inflammation and fibrosis (Fig.4m). Interestingly, the expression of all *Cts* genes was higher in F-C/HF than in M-C/HF. MO reduced the expression level of *Cts* genes in females to similar levels than males. In sum, MO appears to prevent liver injury in female obese offspring by reducing inflammatory processes.

### MO prevents hepatocellular carcinoma development in obese female offspring

Recent research in animal models has elucidated potential programming mechanisms that include altered hepatic function^29-32^ and cellular signaling responses^33,34^. Histological analysis of the liver structure in offspring revealed the presence of marked areas of cell proliferation in female livers. This was in marked contrast to males, where liver biopsies displayed small and scattered proliferative spots. Importantly, MO reduced the cell proliferation areas in females (Fig.5a). By exploring KEGG pathways associated with cancer, we could show that their activity was higher in females than in males, regardless of maternal diet. MO repressed the cell cycle and induced chemical carcinogenesis, notch signaling and retinol metabolism pathways in females. In contrast, MO reduced notch signaling pathway activity in males and induced retinol metabolism (Fig.5b). We extracted all the DEG of the selected cancer pathways (Suppl.TableS4) and isolated two major superfamily genes, namely the UDP-glycosyltransferases (*Ugt)* and the sulfotransferases (*Sult)*, which both showed highly sex-specific expression. Indeed, *Ugt3* and *Ugt2* genes were more highly expressed in males than in females, whereas the expression of *Ugt1* and all *Sult* genes - except for *Sult2a8* - were higher in females (Figs.5c-d). MO induced the expression level of the *Ugt* genes in both sexes (Fig.5c). Remarkably, among the two key genes known as tumor repressors (*Osgin1* and *Stat1*) *Osgin1* was more expressed in M-C/HF than F-C/HF, but both were overexpressed by MO in females (Fig.5e). In line with this, genes promoting cancer development and cell apoptosis (*Ccnd1, Fdps* and *Pik3r1*) had lower expression in M-C/HF than in F-C/HF, and MO reduced the expression in females only (Fig.5f).

**Fig. 5.**
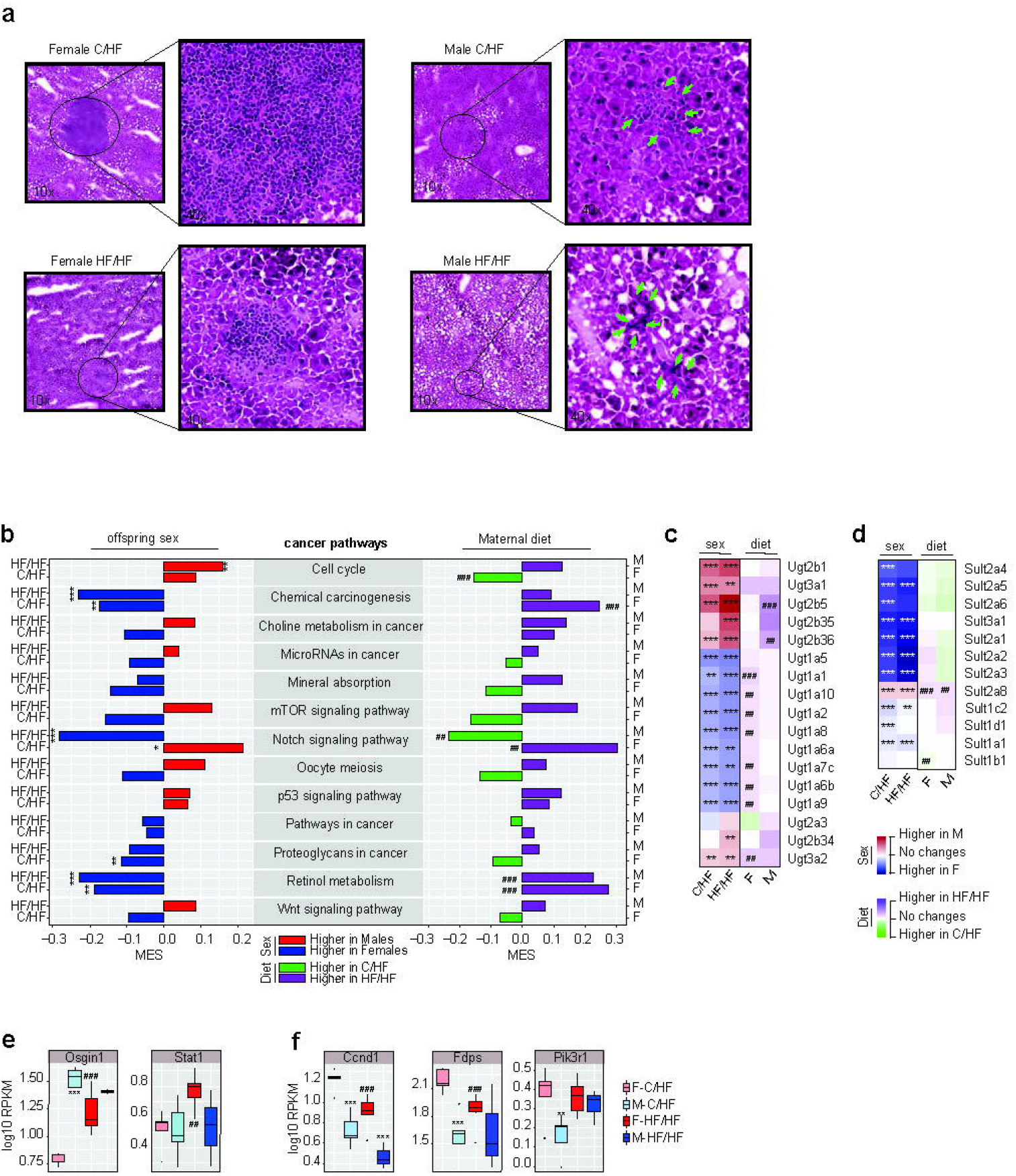
MO has sex-dependent effects on hepatocellular cancer progression in obese offspring. **a** Hematoxylin-eosin (H&E) 10x and 40x images from liver sections of F-C/HF, M-C/HF, F-HF/HF and M-HF/HF; **b** Bar plot presenting the MES between sexes in C/HF and HF/HF and in response to MO in F and M of the KEGG pathways involved in hepatocellular carcinoma. Red and blue bars indicate higher expression in M and F respectively and, green and purple bars indicate higher expression in C/HF and HF/HF groups, respectively. Heatmap of the log2 fold change expression levels of the **c** Ugt - gene family and **d** Sult - gene family; **e-f** Box plots showing expression (RPKM, log10) of genes involved in the nominated cancer pathways. For **a** F-C/HF (*n* =3), M-C/HF (*n* = 3), F-HF/HF (*n* =4) and M-HF/HF (*n* =2). For **b– f** F-C/HF (*n* =5), M-C/HF (*n* = 5), F-HF/HF (*n* =6) and M-HF/HF (*n* = 3). Data are presented as mean ± sem. Differences between two groups (sexes, F versus M; maternal diet C/HF versus HF/HF) were determined by unpaired t-test corrected for multiple comparisons using the Holm–Sidak method, with alpha = 5.000%. For pathway and DEG analysis we used the Benjamini-Hochberg correction with FDR<0.1, when significant. *, M versus F and ^#^, HF/HF versus C/HF, *p* < 0.05; ** or ^##^, *p* < 0.01; *** or ^###^, *p* < 0.001.

Overall, MO tended to reduce cell proliferation markers, and to induce the expression level of tumor repressor and cell apoptosis genes in females, which would indicate a protective mechanism of MO on obese female offspring.

Collectively our data demonstrate that MO modulates differently the metabolism in female and male offspring. We show that sex and MO drive transcriptional and posttranscriptional regulation of major metabolic processes in offspring liver which contribute to the sexual dimorphism in obesity-associated metabolic risks. In figure 6 we summarized the possible mechanisms by which MO may protect female offspring from metabolic impairment, as opposed to male offspring that are impaired. We define differently programmed effects in the female and male liver offspring exposed to MO. Livers from female offspring demonstrate decreased lipogenesis and pro-inflammatory genes, decreased HCC, and remodeling of TG species. These effects were supported by an increased oxidative phosphorylation and browning pathways activity in adipose tissue (Fig.6a). Livers from male offspring show hepatic steatosis, impaired insulin sensitivity and increased inflammation, possibly due to feedback mechanisms from the subcutaneous adipose tissue (Fig.6b).

**Fig. 6.**
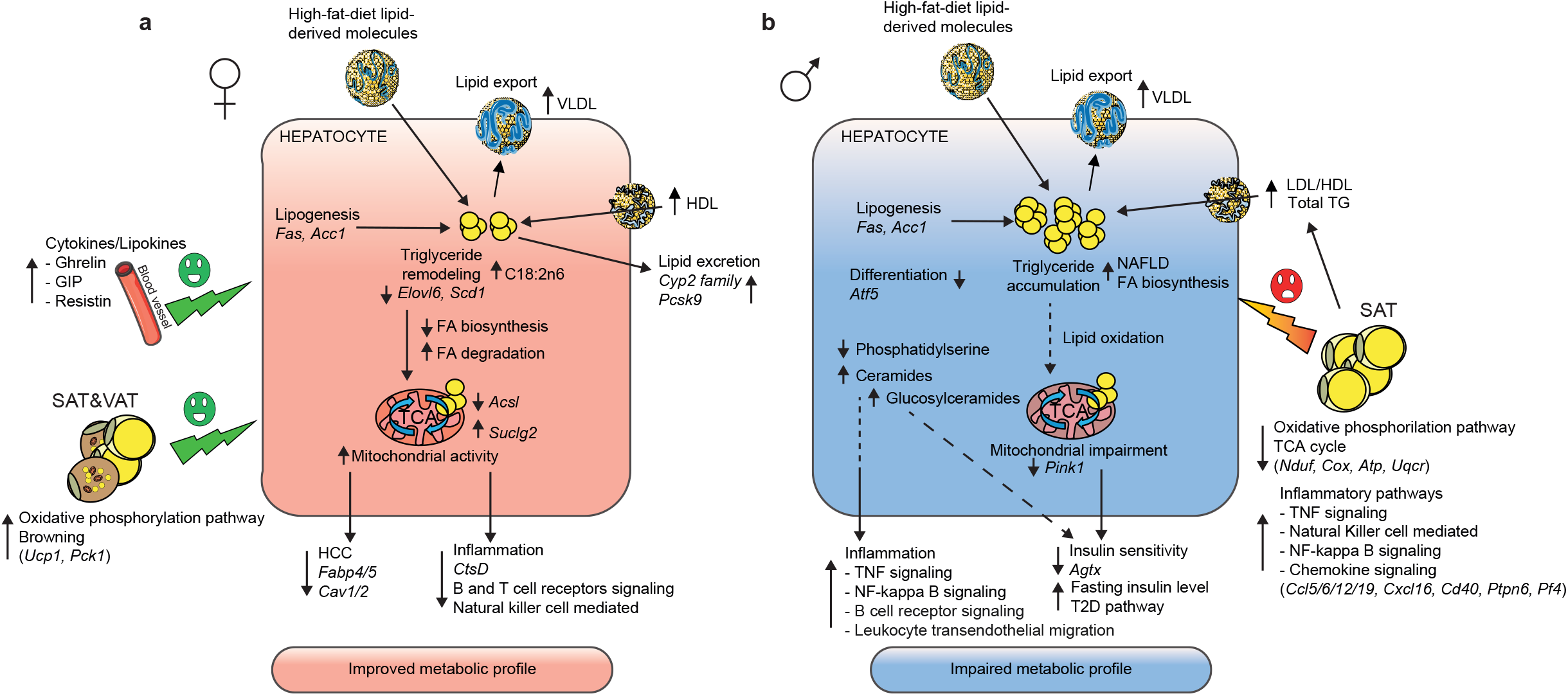
Different programmed metabolic effects in the obese female and male offspring exposed to maternal obesity. **a** Livers from female offspring demonstrate decreased lipogenesis and inflammatory pathways, decreased HCC and remodeling of triglyceride species. Oxidative phosphorylation and browning processes were increased in SAT and VAT, respectively. Circulating cytokine and lipokine levels, ghrelin, GIP and resistin were higher in F-HF/HF compared to F-C/HF and to M-HF/HF. Altogether, these metabolic adaptations may protect the liver from metabolic complications in response to MO; **b** Livers from male offspring demonstrate hepatic steatosis, impaired insulin sensitivity and increased inflammation possibly due to negative feedback signaling from the SAT, including a reduction of oxidative phosphorylation associated with an induction of inflammatory pathways. Arrows represent up-↑ and down-↓ regulation. HCC, hepatocellular carcinoma; SAT, subcutaneous adipose tissue; VAT, visceral adipose tissue; VLDL, very low; LDL, low and HDL, high density lipoproteins.

## DISCUSSION

The current study reveals a number of important mechanisms through which maternal diet primes lipid metabolism differently in obese female and male offspring liver. We previously demonstrated that MO leads to a sexually dimorphic reprogramming of hepatic lipid composition and gene expression, and that this also occurs when the offspring receive a postweaning control diet. We also showed that other organs are affected in a sex-dependent manner^15^. We now show that MO offspring fed an obesogenic diet have sex-specifically altered liver lipidomes explained by accompanying changes in the transcriptomes. When compared to males born to lean mothers, males with obese mothers showed insulin resistance and glucose intolerance at early life stage of life. Moreover, MO did not protect male offspring from the adverse effect of a continued obesogenic diet intake. M-HF/HF had lower body weights early after weaning (3-9 weeks old), while their growth accelerated spontaneously at a later stage (10-15 weeks old). This was associated with metabolic complications including insulin resistance later in life^35^. While M-HF/HF had normal glucose tolerance at MID, they showed impaired glucose tolerance and insulin sensitivity, together with impaired insulin secretion activity and higher circulating level of PAI-1, at END. These findings were opposite to those in female offspring which were not compromised by MO. MO prior to mating and during gestation and lactation did not affect the body weight of obese female offspring, but provoked redistribution of the SAT toward more peripheral and less abdominal accumulation. Interestingly, three major regulators of insulin sensitivity, 3-phosphoinositide-dependent protein kinase-1 (*Pdk1*), NAD(P)H oxidase 4 (*Nox4)* and prolactin receptor (*Prlr*)^36-38^, were significantly influenced by sex. Males showed reduced *Pdk1* and *Prlr* together with elevated *Nox4* expression levels. These genes involved in the development of insulin resistance are controlled by estrogen^39,40^. It is of interest to note, that we previously could demonstrate that feeding a control diet after weaning of male offspring to obese mothers improved insulin sensitivity at END^14^. This indicates that the changes induced by MO *in utero* in male offspring can be reversed by a post-weaning diet.

By using cross-sectional data analysis obtained by multidisciplinary techniques, we showed that lipid profile in the liver of the offspring was sex-dependent, and that it is changed by modulating transcriptional activities, in a sex-dependent manner. We uncovered that relative abundancies of PL, TG and FA lipid species were different between sexes, which may be a key element in the sex-specific metabolic complications in obesity. Most importantly, we confirm our previous findings that MO modulates hepatic TG molecular species in female but not in male offspring^14^. Somewhat surprising, we demonstrated that MO regulates the gene expression in white adipose in male offspring towards an inflammatory pattern, while instead altering it towards more browning and oxidative phosphorylation patterns in females^15^. The mechanisms by which MO can differently modulate epigenetic marks *in utero* between sexes and between tissues remain intriguing and require further investigation.

Desaturases are enzymes that control the balance between saturated, monounsaturated and polyunsaturated FA being incorporated into TG and PL. When fed a control diet after weaning, offspring born from obese mothers showed sex-dependent Δ19 desaturase activity but similar Δ15 desaturase activity^14^. In the current study, male offspring showed lower Δ15 desaturase activity compared to females in both mother diet groups. A low Δ15 is correlated to insulin resistance, abdominal adiposity and predicts the development of type-2 diabetes^41-43^. Long chain acyl co-A synthetases (*Acsl*) are important regulators of FA uptake. *Acsl1* promotes TG accumulation in the liver as opposed to *Acsl3* (localized in lipid droplets) and *Acsl5* (in mitochondria), which regulate lipogenesis and β-oxidation, respectively, thereby being essential for lipid homeostasis^25,44^. Importantly, the *Acsl* isoform expression patterns seen in our study were highly sex- and maternal-diet dependent, and may contribute to the sexual dimorphism in obesity and in responses to MO. Interestingly, *Acsl* expression has been shown to be controlled by estrogen in mammals^45^ and other species^46^.

Alterations of hepatic lipid composition are likely to cause liver damage through various processes, including inflammation, oxidative stress, fibrosis and hepatocellular carcinoma. The expression of cathepsin (Cts) genes was higher in females than in males born to lean mothers, with MO strongly reducing Cts expression in females (to “male” levels). Cts has several functions, including the facilitation of cholesterol excretion and protection against inflammation and CtsD has been identified as a marker of liver inflammation and fibrosis in murine steatohepatitis^47^. In line with this, female offspring of MO showed smaller lipid droplets and reduced cell proliferation and inflammation as compared to those born from lean mothers, which could be an estogen-dependent effect^48,49^. We found sex-specific and maternal diet-dependent changes in the expression of members of the UDP-glucuronosyltransferase (UGT) and sulfotransferase (SULT) gene families. These are essential for the metabolism of xenobiotic and endobiotic substances and may be crucial regulators of hepatic cholesterol and lipid homeostasis^50-53^. The mechanism(s) by which MO and estrogen protect female offspring from liver dysfunction need to be addressed in more extensive future studies.

In conclusion, our detailed studies in mice clearly demonstrate that MO is a preponderant factor for metabolic alterations in offspring. Notably, we show that MO affects hepatic lipid metabolism differently in obese female and male offspring through sex-specific alterations of the expression of genes involved in insulin signaling, liver steatosis, inflammation, fibrosis and carcinoma. A summary of our main findings in female and male offspring is presented in Fig 6. MO can obviously modulate gene expression between sexes as well as between tissues. Interestingly, while MO clearly has negative metabolic effects in male offspring, it seems to protect from the development of insulin resistance and even liver fibrosis and carcinoma in female offspring. The identification of several sex- and maternal diet-regulated genes involved in these processes should now permit further exploration of their possible use to target cardiometabolic risk in humans.

## Methods

### Mice and diet

All animal procedures were approved by the local Ethical Committee of the Swedish National Board of Animal Experiments. Virgin C57Bl6/J female dams and male sires were received at 4 weeks of age. F0 dams were housed in pairs in six different cages and fed either the control diet (C; D12450H, Research Diets, NJ, USA; 10% kcal fat from soybean oil and lard; n=6, F0-CD) or the high fat diet (HF; D12451, Research Diets, NJ, USA; 45% kcal fat from soybean oil and lard; n=6, F0-HFD) for six weeks before mating. Sires remained on control diet (C) until sacrifice. After six weeks of their respective diet two F0 dams were mated with one F0 sire. During this short mating period (up to five days) sires were on the same HF as dams in the group (experimental unit). The sires spermatozoa were unlikely affected by the HFD given a general sperm maturation time of approximately 35 days ^54^. After mating, F0 males and pregnant dams were separated. F0 dams were continuously exposed to their respective diets throughout pregnancy and until the end of the lactation period. The F1 offspring were weaned at 3-week of age. Afterwards, F1 males and females were sex-separated, three to five animals were housed per cage and fed with HFD until the end of the study (Fig.1a). To simplify the naming convention, the group of offspring born from HFD fed dams were named HF/HF (for HFD F0 dam and HFD F1 offspring) and the group of offspring born from CD fed mother named C/HF (for CD mother and HFD offspring). All mice were housed in a 23°C temperature-controlled 12h light/dark room, with free access to water and food unless specified. Body weight was recorded weekly throughout the study in all groups. Average food intake in offspring was recorded twice a week for three weeks in four different cages containing grouped mice (n=3-5 animals per cage) around 4-month of age and at least one week after recovering of *in vivo* experiments. We then calculated the average food intake per cage during the three experimental weeks. We reported it to the average food intake per mouse according to the number of animals in the cage.

### *In vivo* magnetic resonance imaging (MRI)

Animals were anesthetized using isoflurane (4% for sleep induction and ∼2% for sleep maintenance) in a 3:7 mixture of oxygen and air, before being positioned prone in the MR-compatible animal holder. Respiration was monitored during scanning (SA-instruments, Stony Brook, NY, USA). Core body temperature was maintained at 37°C during scanning using a warm air system (SA-instruments, Stony Brook, NY, USA). Magnetic resonance imaging (MRI) images (n=5-7 per group) were collected using a 9.4 T horizontal bore magnet (Varian Yarnton UK) equipped with a 40 mm millipede coil, as previously detailed^5^. Fiji software (http://fiji.sc) was used to compute the volume of fat in different regions of interest in the body. Visceral fat (VAT) was calculated as the difference between the total (TF) signal and the total subcutaneous fat (Total SAT) signal in the abdominal region. Abdominal fat (ABD) comprises the SAT and the VAT fat signals from the abdominal region and the SAT in ABD was calculated as the difference between the ABD fat and the VAT. MRI experiments were performed on the same mouse (F1) at the age of 3 months (MID) and 6 months (END).

### *In vivo* localized proton magnetic resonance spectra (^1^H-MRS)

As for the MRI scanning, animals were anesthetized using isoflurane, respiration was monitored, and core body temperature maintained at 37°C during scanning. In addition, heart beats were recorded using an electrocardiogram system. Localized proton magnetic resonance spectra (^1^H-MRS) from the liver (n=5-7 per group) were acquired from a 2×2×2 mm^3^ voxel localized in the left lobe with excitation synchronized to the first R-wave within the expiration period, as detailed^55,56^. Spectroscopy data were processed using the LCModel analysis software (http://s-provencher.com/pub/LCModel/manual/manual.pdf). “Liver 9” was used as a base with all signals occurring in the spectral range of 0 to 7 ppm (water resonance at 4.7ppm) simulated in LCModel. All concentrations were derived from the area of the resonance peaks of the individual metabolites. Only the fitting results with an estimated standard deviation of less than 20% were further analyzed. ^1^H-MRS spectra revealed nine lipid signals (peaks) in the mouse liver. Peak assignments were based on published data^55,56^. As for the MRI, ^1^H-MRS experiments were repeated twice on the same animal at MID and END.

### *In vivo* metabolic tolerance tests

At MID and END, F1 mice were fasted for 6h prior to the oral glucose tolerance test (OGTT) and for 4h prior to the insulin tolerance test (ITT), both performed as detailed^57^. Briefly, at time zero (T0) peripheral glucose level was measured at the tail using a One-Touch ultra-glucometer (AccuChek Sensor, Roche Diagnostics) and at T15, T30, T60 and T120 min. For the OGTT, extra blood was collected at each time-point and later plasma was separated by centrifugation (15min at 2,000 RPM) and stored at −80 °C for insulin measurement using a Rat/Mouse Insulin Elisa kit (EMD Millipore - EZRMI13K). Prior to sacrifice, mice were fasted for 2h and anesthetized with 4% isoflurane. Blood glucose level was measured with a OneTouch Ultra glucometer (AccuChek Sensor, Roche Diagnostics). Subsequently, mice were exsanguinated via cardiac puncture and blood saved for plasma analysis. The whole liver was quickly removed and washed into PBS. Several pieces of left lobe of the liver were collected, fresh-frozen into liquid nitrogen and stored at - 80°C until further analysis.

### Liver histology

For hematoxylin and eosin (H&E) staining, the livers were frozen in OCT embedding matrix and on dry ice. Sectioning and staining were done according to standard histological procedures.

### Biochemical analysis of plasma

Within 15 min after blood collection, plasma was separated by centrifugation (15min at 2,000 RPM). Plasma total triglycerides (Total TG) and total cholesterol (Total Chol) were measured by enzymatic assay using commercially available kits (Roche Diagnostics GmbH, Mannheim and mti Diagnostic GmbH, Idstein, Germany). Cholesterol lipoprotein fractions in serum were determined as described^58^. Briefly, sera from each individual mouse were separated by size exclusion chromatography using a Superose and PC 3.2/30 column (Pharmacia Biotech, Uppsala, Sweden). Reagent (Roche Diagnostic, Mannheim, Germany) was directly infused into the eluate online and the absorbance was measured. The concentration of the different lipoprotein fractions was calculated from the area under the curves of the elution profiles by using the EZChrom Elite software (Scientific Software; Agilent Technologies, Santa Clara, CA).

### Immunoassay for adipokine levels

Within 15 min after blood collection, plasma was separated by centrifugation (15min at 2,000 RPM). A Multiplexed bead immunoassay was used to measure adipokine levels using a commercially available kit (Bio-Plex Pro Mouse Diabetes 8-Plex Assay #171F7001M) according to manufacturer’s instructions.

### Lipidomic

#### Fatty acid analysis using gas chromatography – mass spectrometry (GC-MS)

Total lipid extracts were obtained using a modified Bligh and Dyer method ^59^ and after transmethylation, the fatty acids were analyzed by gas chromatography followed by mass spectrometry (GC-MS) ^60,61^. Aliquots of the lipid extracts corresponding to 2.5 μg of total phospholipid, were transferred into glass tubes and dried under a nitrogen stream. Resulting lipid films were dissolved in 1 mL of *n*-hexane containing a C19:0 as internal standard (1.03 μg mL^−1^, CAS number 1731-94-8, Merck, Darmstadt, Germany) with addition of 200 μL of a solution of potassium hydroxide (KOH, 2 M) in methanol, followed by 2 min vortex. Then, 2 mL of a saturated solution of sodium chloride (NaCl) was added, and the resulting mixture was centrifuged for 5 minutes at 626 x g for phase separation. Cholesterol was removed from the organic phase according to the Lipid Web protocol (https://lipidhome.co.uk/ms/basics/msmeprep/index.htm). A 1 cm silica column in a pipette tip with wool was pre-conditioned with 5 mL of hexane (high-performance liquid chromatography (HPLC) grade). Methyl esters were added to the top of the tip and recovered by elution with hexane:diethyl ether (95:5, v/v, 3 mL), and thereafter dried under a nitrogen current. Fatty acid methyl esters (FAMEs) were dissolved in 100 µL, and 2.0 μL were injected in GC-MS (Agilent Technologies 8860 GC System, Santa Clara, CA, USA). GC-MS was equipped with a DB-FFAP column (30m long, 0.32 mm internal diameter, and 0.25 μm film thickness (J & W Scientific, Folsom, CA, USA)). The GC equipment was connected to an Agilent 5977B Mass Selective Detector operating with an electron impact mode at 70 eV and scanning the range *m/z* 50–550 in a 1 s cycle in a full scan mode acquisition. Oven temperature was programmed from an initial temperature of 58°C for 2 min, a linear increase to 160°C at 25°C min^−1^, followed by linear increase at 2°C min^−1^ to 210 °C, then at 20 °C min^−1^ to 225°C, standing at 225°C for 20 min. Injector and detector temperatures were set to 220 and 230°C, respectively. Helium was used as the carrier gas at a flow rate of 1.4 mL min^−1^. GCMS5977B/Enhanced Mass Hunter software was used for data acquisition. To identify fatty acids (FA), the acquired data were analysed using the qualitative data analysis software Agilent MassHunter Qualitative Analysis 10.0. FA identification was performed by MS spectrum comparison with the chemical database NIST library and confirmed with the literature.

The total ω-3 content was calculated as the summed total of ω-3 PUFA of C18:3ω-3, C20:5ω-3, C22:5ω-3 and C22:6ω-3. Total ω-6 content was calculated as the summed total of C18:2ω-6, C18:3ω-6, C20:2ω-6, C20:3ω-6 and C20:4ω-6 contents. Total ω-9 MUFA were calculated as the summed of C16:1ω-9 and C18:1ω-9 contents. Total ω-11 MUFA were calculated as the summed of C16:1ω-11 and C18:1ω-11 contents.

#### Phospholipids (PL), sphingolipids (SL) and triglycerides (TG) analysis by Liquid Chromatography - Mass Spectrometry

Total lipid extracts from the left lobe of the liver were separated using a HPLC system (Ultimate 3000 Dionex, Thermo Fisher Scientific, Bremen, Germany) with an autosampler coupled online to a Q-Exactive hybrid quadrupole Orbitrap mass spectrometer (Thermo Fisher Scientific, Bremen, Germany), adapted from^59,62^. Briefly, the solvent system consisted of two mobile phases: mobile phase A (ACN/MeOH/water 50:25:25 (v/v/v) with 2.5 mM ammonium acetate) and mobile phase B (ACN/MeOH 60/40 (v/v) with 2.5 mM ammonium acetate). Initially, 10% of mobile phase A was held isocratically for 2 min, followed by a linear increase to 90% of A within 13 min and a maintenance period of 2 min, returning to the initial conditions in 3 min, followed by a re-equilibration period of 10 min prior to the next injection. Five μg of phospholipid (PL) from total lipid extracts were mixed with 4 μL of phospholipid standard mixture (dMPC - 0.02 µg, dMPE - 0.02 µg, SM - 0.02 µg, LPC - 0.02 µg, TMCL - 0.08 µg, dPPI - 0.08 µg, dMPG - 0.012 µg, dMPS - 0.04 µg, Cer - 0.04 µg, dMPA - 0.08 µg) and 91 μL of solvent system (90% of eluent B and 10% of eluent A). Five μL of each dilution were introduced into the AscentisSi column (10 cm × 1 mm, 3 μm, Sigma-Aldrich, Darmstadt, Germany) with a flow rate of 50 μL min^−1^. The temperature of the column oven was maintained at 35 °C. The mass spectrometer with Orbitrap technology operated in positive (electrospray voltage 3.0 kV) and negative (electrospray voltage −2.7 kV) ion modes with a capillary temperature of 250 °C, a sheath gas flow of 15 U, a high resolution of 70 000 and AGC target 1e6. In MS/MS experiments, cycles consisted of one full scan mass spectrum and ten data-dependent MS/MS scans (resolution of 17 500 and AGC target of 1e5), acquired in each polarity. Cycles were repeated continuously throughout the experiments with the dynamic exclusion of 60 s and an intensity threshold of 2e4. Normalized collisional energy ranged between 20, 25, and 30 eV.

#### Reagents/Chemicals for LC-MS analysis

Phospholipid internal standards 1,2-dimyristoyl-*sn*-glycero-3-phosphocholine (dMPC), 1-nonadecanoyl-2-hydroxy-*sn*-glycero-3-phosphocholine (LPC), 1,2-dimyristoyl-*sn*-glycero-3-phosphoethanolamine (dMPE), N-palmitoyl-D-*erythro*-sphingosylphosphorylcholine (NPSM – SM d18:1/17:0), N-heptadecanoyl-D-erythro-sphingosine (Cer d18:1/17:0), 1,2-dimyristoyl-*sn*-glycero-3-phospho-(10-rac-)glycerol (dMPG), 1,2-dimyristoyl-*sn*-glycero-3-phospho-L-serine (dMPS), tetramyristoylcardiolipin (TMCL), 1,2-dimyristoyl-*sn*-glycero-3-phosphate (dMPA) and 1,2-dipalmitoyl-*sn*-glycero-3-phosphatidylinositol (dPPI) were purchased from Avanti Polar Lipids, Inc. (Alabaster, AL). HPLC grade dichloromethane, methanol and acetonitrile were purchased from Fisher scientific (Leicestershire, UK). All the reagents and chemicals used were of the highest grade of purity commercially available and were used without further purification. The water was of Milli-Q purity (Synergy®, Millipore Corporation, Billerica, MA).

Spectra were analyzed in positive and negative mode, depending on the lipid class. Ceramides (Cer), glucosylceramides (GlcCer), phosphatidylethanolamine (PE), lyso phosphatidylethanolamine (LPE), phosphatidylcholine (PC), lysophosphatidylcholine (LPC), and sphingomyelin (SM) were analyzed in the LC-MS spectra in the positive ion mode, and identified as [M+H]^+^ ions, while cardiolipin (CL), phosphatidylserine (PS), phosphatidylinositol (PI), lysophosphatidylinositol (LPI) and phosphatidylglycerol (PG) species were analyzed in negative ion mode, and identified as [M−H]^−^ ions. Molecular species of triacylglycerol (TG) were also analyzed in positive ion mode as [M+NH_4_]^+^ ions. Data acquisition was carried out using the Xcalibur data system (V3.3, Thermo Fisher Scientific, USA). The mass spectra were processed and integrated through the MZmine software (v2.32)^63^. This software allows for filtering and smoothing, peak detection, alignment and integration, and assignment against an in-house database, which contains information on the exact mass and retention time for each PL, Cer and TG molecular species. During the processing of the data by MZmine, only the peaks with raw intensity higher than 1e4 and within 5 ppm deviation from the lipid exact mass were considered. The identification of each lipid species was validated by analysis of the LC-MS/MS spectra. The product ion at *m/z* 184.07 (C_5_H_15_NO_4_P), corresponding to phosphocholine polar head group, observed in the MS/MS spectra of the [M+H]^+^ ions allowed to pinpoint the structural features of PC, LPC and SM molecular species under MS/MS conditions^59^, which were further differentiated based on *m/z* values of precursor ions and characteristic retention times. PE and LPE molecular species ([M+H]^+^ ions) were identified by MS/MS based on the typical neutral loss of 141 Da (C_2_H_8_NO_4_P), corresponding to phosphoethanolamine polar head group. These two classes were also differentiated based on *m/z* values of precursor ions and characteristic retention times. The [M+H]^+^ ions of Cer and GlcCer molecular species were identified by the presence of product ions of sphingosine backbone in MS/MS spectra, such as ions at *m/z* 264.27 (C_18_H_34_N) and 282.28 (C_18_H_36_NO) for sphingosine d18:1^64^, together with the information on *m/z* values of precursor ions and characteristic retention times. The PG molecular species were identified by the [M−H]^−^ ions and based on the product ions identified in the corresponding MS/MS spectra, namely the product ions at *m/z* 152.99 (C_3_H_6_O_5_P) and 171.01 (C_3_H_8_O_6_P). PI and LPI, also identified as [M-H]^-^ ions, were confirmed the product ions at *m/z* 223.00 (C_6_H_8_O_7_P), 241.01 (C_6_H_10_O_8_P), 297.04 (C_9_H_14_O_9_P) and 315.05 (C_9_H_16_O_10_P), which all derived from phosphoinositol polar head group^59,65^. The [M−H]^-^ ions of PS molecular species were identified based on product ions at *m/z* 152.99 (C_3_H_6_O_5_P) in MS/MS spectra, retention time and *m/z* values of precursor ions. CL molecular species ([M−H]^−^ ions) were characterized by MS/MS with identification of ions at *m/z* 152.99 (C_3_H_6_O_5_P), carboxylate anions of fatty acyl chains (RCOO^−^), product ions corresponding to phosphatidic acid anion and phosphatidic acid anion plus 136 Da as previously reported^65^. Negative ion mode MS/MS data were used to identify the fatty acid carboxylate anions RCOO^−^, which allowed the assignment of the fatty acyl chains esterified to the PL precursor. The MS/MS spectra of [M+NH_4_]^+^ ions of TGs allowed the assignment of the fatty acyl substituents on the glycerol backbone^66^.

### Unsupervised clustering

The raw data matrix of the lipid spectra was distributed column-wise by sample IDs and row-wise by PL names. The TMM method was used to normalize between samples^67^. Unsupervised clustering was then performed using the principal component analysis (PCA) plot option in R. The PCA plot is based on the two most variant dimensions in which the PL parameters with duplicated data are filtered out.

### RNA isolation, purity and integrity determination

Liver, SAT and VAT total RNA was extracted using QIAGEN miRNeasy Mini Kit (217004, Qiazol). RNA concentration was measured by nanodrop®. RNA was treated with RNase-free DNase (79254) according to the manufacturer’s instructions. cDNA libraries were prepared for bulk-RNA sequencing as previously detailed^15^.

### Bulk RNA-seq mapping

All raw sequence reads available in FastQ format were mapped to the mouse genome (mm10) using Tophat2 with Bowtie2 option ^68,69^, as described previously^15^. Raw read counts for each gene were calculated using featureCounts from the subread package^70^.

### Bulk RNA-seq differential gene expression analysis

A differential gene expression analysis was performed using DEseq2^71^. The differentially expressed genes (DEG) were identified by adjusted *p*-value for multiple testing using Benjamini-Hochberg correction with False Discovery Rate (FDR) values less than 0.1.

### Pathway analysis

A Gene Set Enrichment Analysis (GSEA) was performed using the KEGG pathways dataset. Genes were ranked in descending order according to the log_2_ fold change (log_2_FC) of expression. Differences between the ranks of genes in a pathway were compared to other genes. For each queried pathway,

if gene *i* is a member of the pathway, it is defined as:

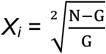

if gene *i* is not a member of the pathway, it is defined as:

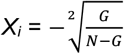

here *N* is the total number of genes and *G* indicates the number of genes in the query pathway. Next, a max running sum across all *N* genes Maximum Estimate Score (MES) is calculated as:

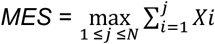

The permutation test was performed with 1000 times to judge the significance of MES values. The queried pathway with a nominal *p*-value less than 0.05 and FDR values less than 0.1 are considered to be significantly enriched. The positive MES value indicates enrichment (up-regulation) whereas a negative MES value indicates depletion (down-regulation) of a pathway activity.

### Gene Ontology (GO) enrichment analysis

Gene Ontology enrichment analysis is performed using online software AmiGO website (http://amigo.geneontology.org/amigo), where the significant enrichment GO terms was identified using Fisher’s Exact test with P values ≤ 0.05.

### Statistical analyses

Data are expressed as mean ± sem. Differences between the four group (female and male offspring sex and C and HF diet mother groups (F-C/HF, M-C/HF, F-HF/HF and M-HF/HF) were determined using two-way ANOVA with diet (D) and sex (S) as independent variables, followed by Tukey’s multiple comparison post hoc test when significant (p<0.05).

Differences between two groups (sexes, F *versus* M; maternal diet C/HF *versus* HF/HF) were determined by t-test corrected for multiple comparisons using the Holm-Sidak method, with alpha=5.000%. *, p<0.05 M *vs* F and ^#^, p<0.05 HF/HF *vs* C/HF were considered significant. ** or ^##^, p<0.01; *** or ^###^, p<0.001.

## Acknowledgements

The MRI and MRS experiments were performed at the Department of Comparative Medicine/Karolinska Experimental Research and Imaging Centre at Karolinska University Hospital, Solna, Sweden. We thank Peter Damberg and Sahar Nikkhou Aski for excellent assistance to develop the sequence for proton-magnetic resonance spectroscopy in the liver. We thank Ingela Arvidsson for excellent help at the FLPC for lipoprotein profiling. We thank Byambajav Buyandelger, Sonja Gustafsson, Jianping Liu from the single cell facility, Karolinska institute in Huddinge for excellent assistance for the SmartSeq2 experiment. This work and M.K.A. were supported by the Novo Nordisk Foundation (NNF14OC0010705), by the Lisa and Johan Grönbergs Foundation (2019-00173) and by AstraZeneca (ICMC). L.A.H. is supported by grants from FCT - Fundação para a Ciência e a Tecnologia (UID/BIM/04501/2020), CCDRC (CENTRO-01-0145-FEDER-000003) and CCDRC (CENTRO-01-0246-FEDER-000018), M.R.D., D.C. and T.M. are supported by CESAM (UIDP/50017/2020+UIDB/50017/2020) and LAQV/REQUIMTE (UIDB/50006/2020). Fetus in Fig.1a was created by Servier Medical Art “newborn mouse.” In Fig.6, lipoprotein cells were designed using Servier Medical Art “lipids” http://smart.servier.com/. Open Access licensed under a Creative Common Attribution 3.0 Generic License https://creativecommons.org/licenses/by/3.0/legalcode.

## Author Contributions

M.K.A. conceptualized and designed the study. C.S., M.G.G and M.K.A. performed animal experiments; C.S. and M.K.A. collected and analyzed all generated data; L.H., D.C., T.M. and M.R.D. performed the lipidomics and wrote the method for lipidomic; C.S. performed RNA sequencing experiments; C.S. and X.L. performed the bioinformatics; C.S. and M.K.A. designed the figures and wrote the manuscript; B.A. and C.K. substantially participated to the manuscript review. The manuscript was edited and approved by all authors. M.K.A is the guarantor of this work and, as such, had full access to all the data in the study and takes responsibility for the integrity of the data and the accuracy of the data analysis. All authors approved the final version of the manuscript.

## Data availability

The raw data generated for lipidomics and RNA sequencing are available as described below For VAT and SAT RNA sequencing, SRA data: PRJNA662930

For Liver RNA sequencing, SRA data: PRJNA723771

For the lipidomic, SRA data: https://figshare.com/s/ac91b57eaa0f5c560d3d

## Figure legends

**Supplementary Figure S1. Metabolic pathways in liver of offspring. a** DEG and differential pathway activity in response to MO obtained from Smart-seq2 data analysis in subcutaneous (SAT) and visceral (VAT) adipose tissues of male and female offspring; **b** Clustered heatmap of all genes obtained from Smart-seq2 in liver between sexes in C/HF and HF/HF (sex) and in response to MO in F and M (diet); **c** Clustered heatmap of all KEGG pathway enrichment analysis presenting the MES levels between sexes in C/HF and HF/HF (sex) and in response to MO in F and M (diet). Data are presented as mean ± sem. F-C/HF (*n* =5), M-C/HF (*n* = 5), F-HF/HF (*n* =6) and M-HF/HF (*n* = 3).

**Supplementary Figure S2. Relative abundance of TG species in offspring’s liver detected by LC-MS**. Relative abundance of **a** Low abundant short TG; **b** Low abundant long TG and **c** High abundant TG species detected by LC-MS in F-C/HF (red open bars), M-C/HF (blue open bars), F-HF/HF (red stripped bars) and M-HF/HF (blue stripped bars). For **a-c** F-C/HF (*n* =4), M-C/HF (*n* = 4), F-HF/HF (*n* =4) and M-HF/HF (*n* = 3). Data are presented as mean ± sem. Differences between two groups (sexes, F versus M; maternal diet C/HF versus HF/HF) were determined by unpaired t-test corrected for multiple comparisons using the Holm–Sidak method, with alpha = 5.000%. *, M versus F and ^#^, HF/HF versus C/HF, *p* < 0.05; ** or ^##^, *p* < 0.01; *** or ^###^, *p* < 0.001.

**Supplementary Figure S3. Sex and MO-dependent relative abundance of fatty acid (FA) species in offspring. a** Relative abundance of fatty acid (FA) species contained into the total TG and PL in F-C/HF (red open bars), M-C/HF (blue open bars), F-HF/HF (red stripped bars) and M-HF/HF (blue stripped bars); Pie charts of the **b** ω-3, ω-6, ω-9 and ω-11 FA synthesis pathways and **c** FA saturation profile in F-C/HF, M-C/HF, F-HF/HF and M-HF/HF; **d** Delta 9 and delta 5 desaturase activity in F-C/HF (red open box), M-C/HF (blue open box), F-HF/HF (red stripped box) and M-HF/HF (blue stripped box). For **a-d** F-C/HF (*n* =4), M-C/HF (*n* = 4), F-HF/HF (*n* =4) and M-HF/HF (*n* = 3). Data are presented as mean ± sem. Differences between two groups (sexes, F versus M; maternal diet, C/HF versus HF/HF) were determined by unpaired t-test corrected for multiple comparisons using the Holm–Sidak method, with alpha = 5.000%. *, M versus F and ^#^, HF/HF versus C/HF, *p* < 0.05; ** or ^##^, *p* < 0.01; *** or ^###^, *p* < 0.001.

**Supplementary Figure S4. PL lipid species relative abundance in liver**. Relative hepatic levels of **a** PC and LPC; **b** PE and LPE lipid classes; Low, moderate and high relative levels of **c** PC and **d** PE species; Low and high relative levels of **e** LPC and **f** LPE species in F-C/HF (red open bars), M-C/HF (blue open bars), F-HF/HF (red stripped bars) and M-HF/HF (blue stripped bars). For **a-f** F-C/HF (*n* =4), M-C/HF (*n* = 4), F-HF/HF (*n* =4) and M-HF/HF (*n* = 3). Data are presented as mean ± sem. Two-way ANOVA (sex (S), mother diet (D), interaction (I) between sex and diet, and (ns) for not significant) followed by Tukey’s multiple comparisons test when significant (*p* < 0.05). Differences between two groups (sexes, F versus M; maternal diet, C/HF versus HF/HF) were determined by unpaired t-test corrected for multiple comparisons using the Holm–Sidak method, with alpha = 5.000%. *, M versus F and ^#^, HF/HF versus C/HF, *p* < 0.05; ** or ^##^, *p* < 0.01; *** or ^###^, *p* < 0.001.

**Supplementary Figure S5. Relative abundance of PL lipid classes and species**. Relative hepatic levels of **a** PS, PG, CL, PI, Cer and SM lipid classes; Low and high relative levels of **b** PS; **c** PG; **d** CL; **e** PI; **f** Cer; and **g** SM species in C/HF (open bars) and HF/HF (stripped bars) F (red bars) and M (blue bars). Data are presented as mean ± sem. Two-way ANOVA (sex (S), mother diet (D), interaction (I) between sex and diet, and (ns) for not significant) followed by Tukey’s multiple comparisons test when significant (*p* < 0.05). Differences between two groups (sexes, F versus M; maternal diet, C/HF versus HF/HF) were determined by unpaired t-test corrected for multiple comparisons using the Holm–Sidak method, with alpha = 5.000%. *, M versus F and ^#^, HF/HF versus C/HF, *p* < 0.05; ** or ^##^, *p* < 0.01; *** or ^###^, *p* < 0.001.

**Supplementary Figure S6. Expression levels of genes involved in hepatic lipid and energy metabolism. a** Box plots showing the expression (RPKM, log10) of genes of the lipid and energy metabolism pathways. F-C/HF (*n* =5), M-C/HF (*n* = 5), F-HF/HF (*n* =6) and M-HF/HF (*n* = 3). Data are presented as mean ± sem. For analysis we used the Benjamini-Hochberg correction with FDR<0.1, when significant. *, M versus F and ^#^, HF/HF versus C/HF, *p* < 0.05; ** or ^##^, *p* < 0.01; *** or ^###^, *p* < 0.001.

