## Supplementary Files for "Molecular programming *in utero* modulates hepatic lipid metabolism and adult metabolic risk in obese mother offspring in a sex-specific manner"

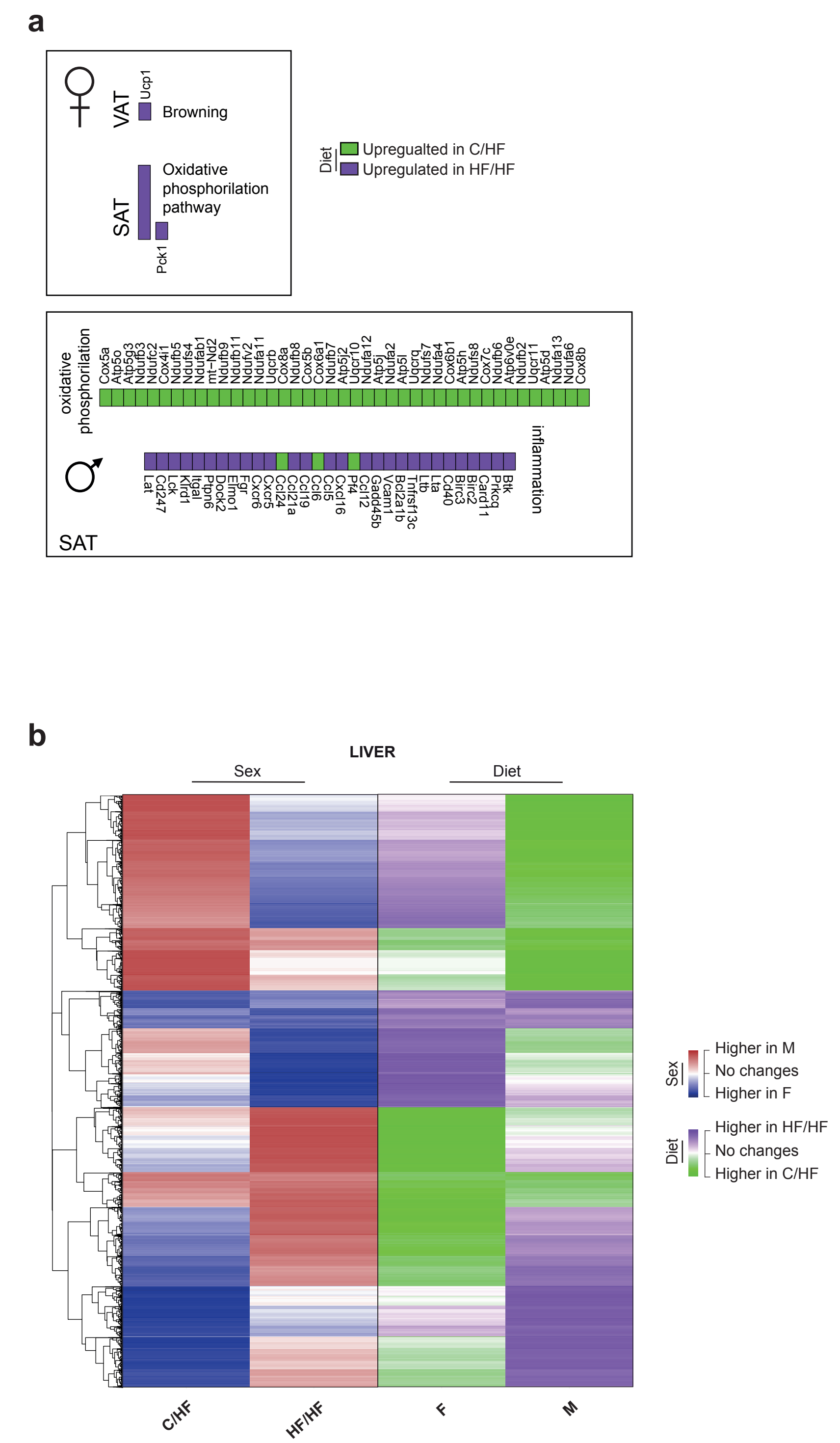


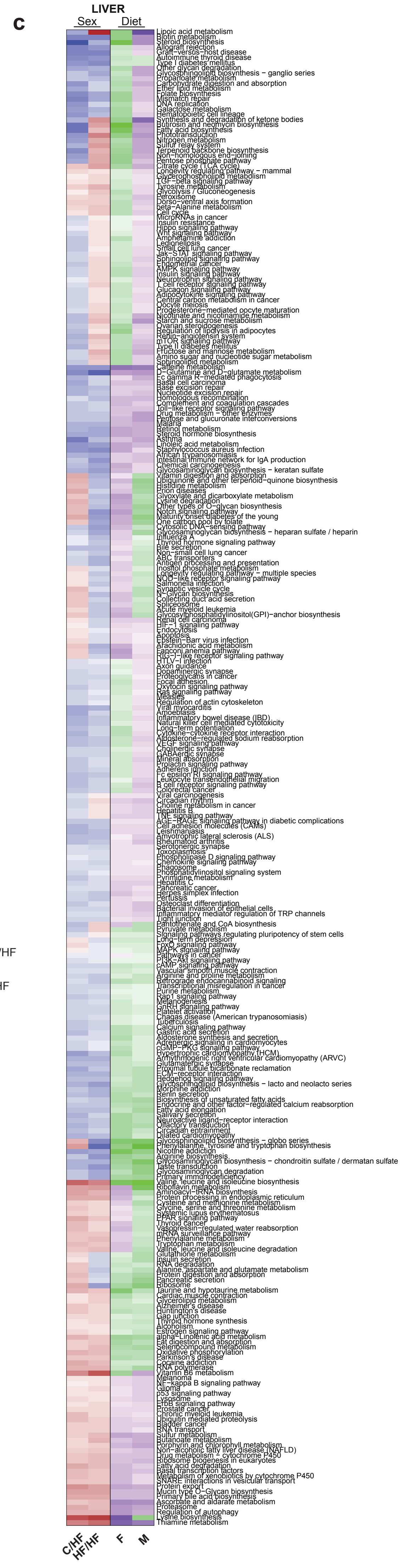


Supplementary Figure S1

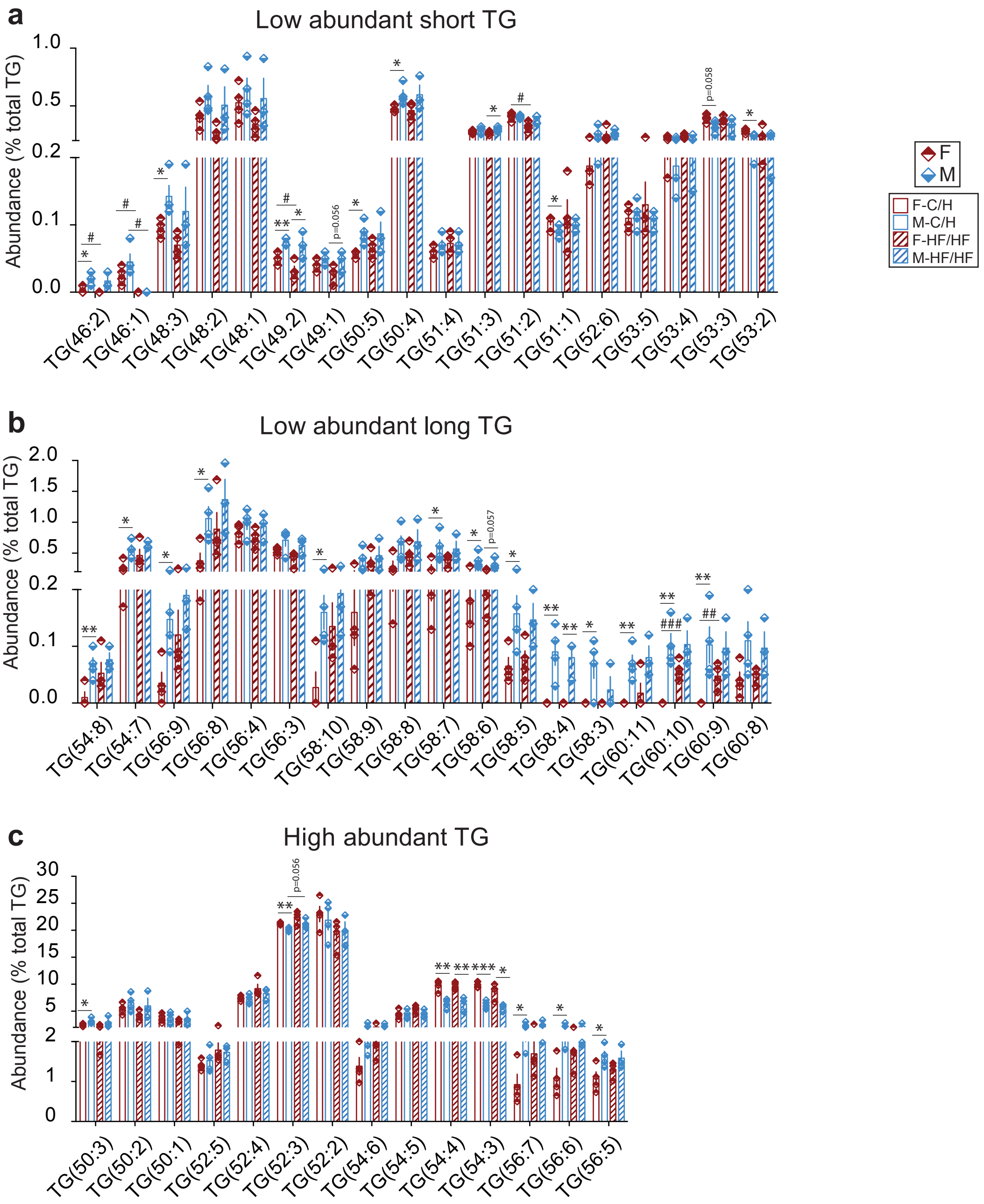


Supplementary Figure S2

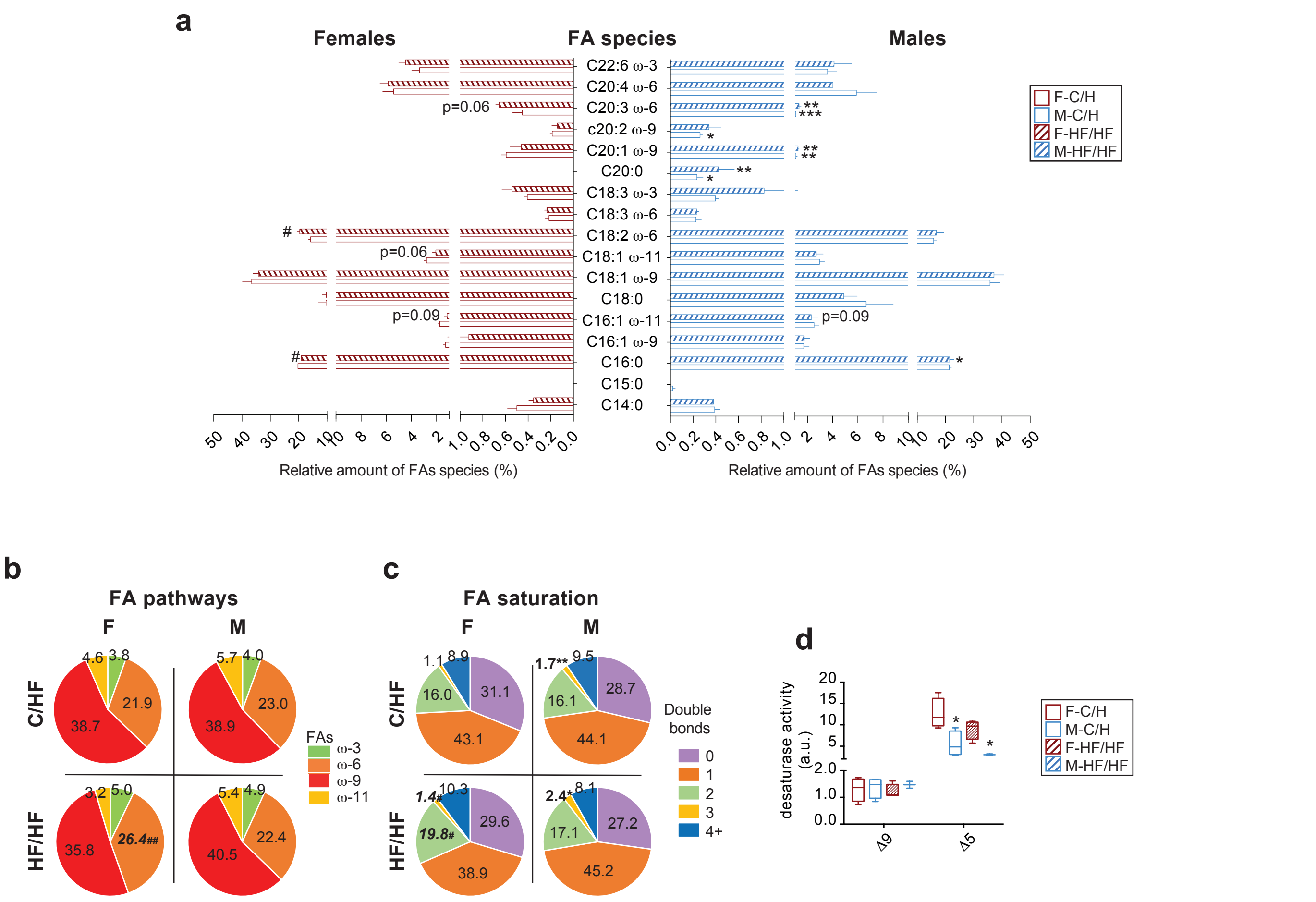


Supplementary Figure S3

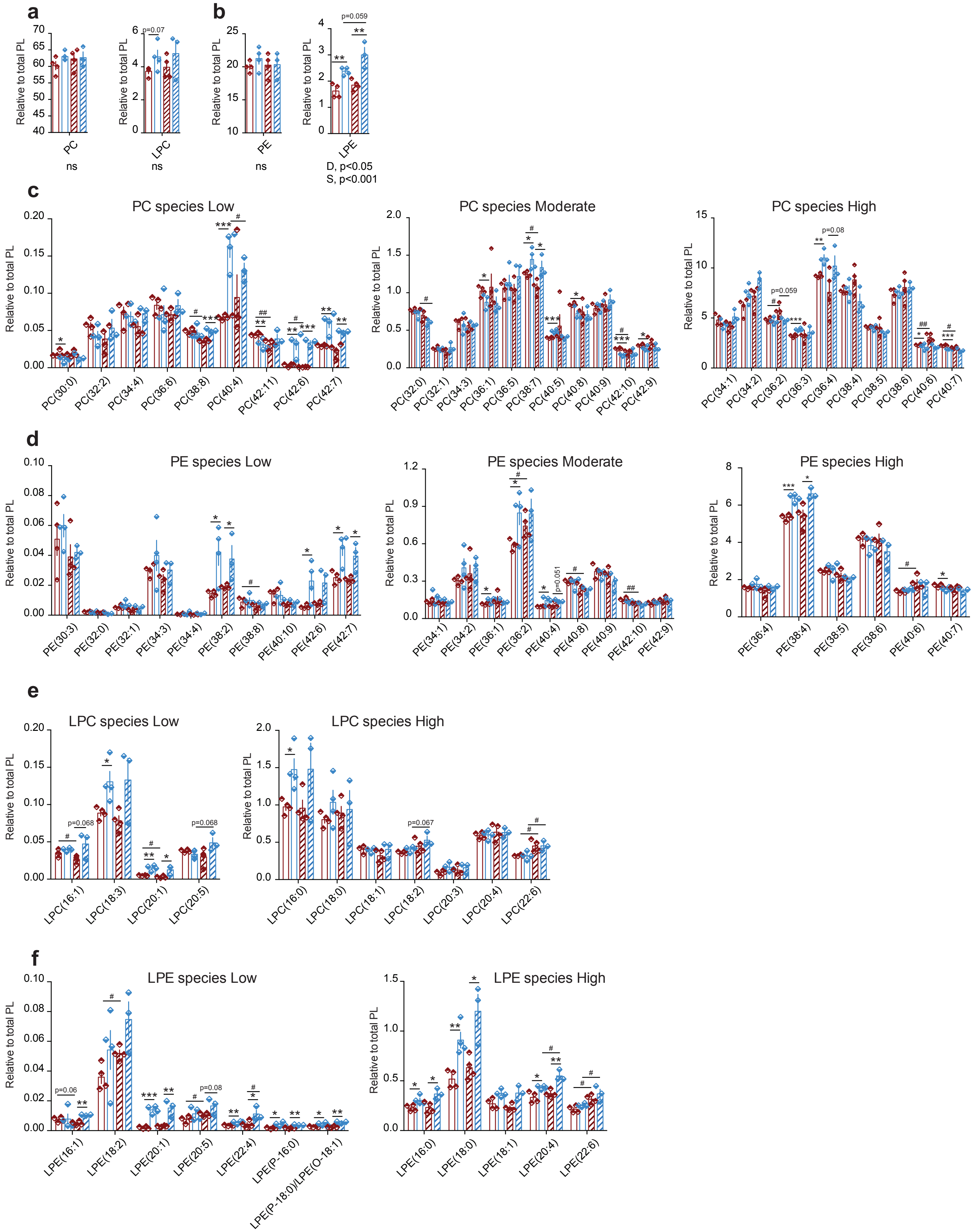


Supplementary Figure S4

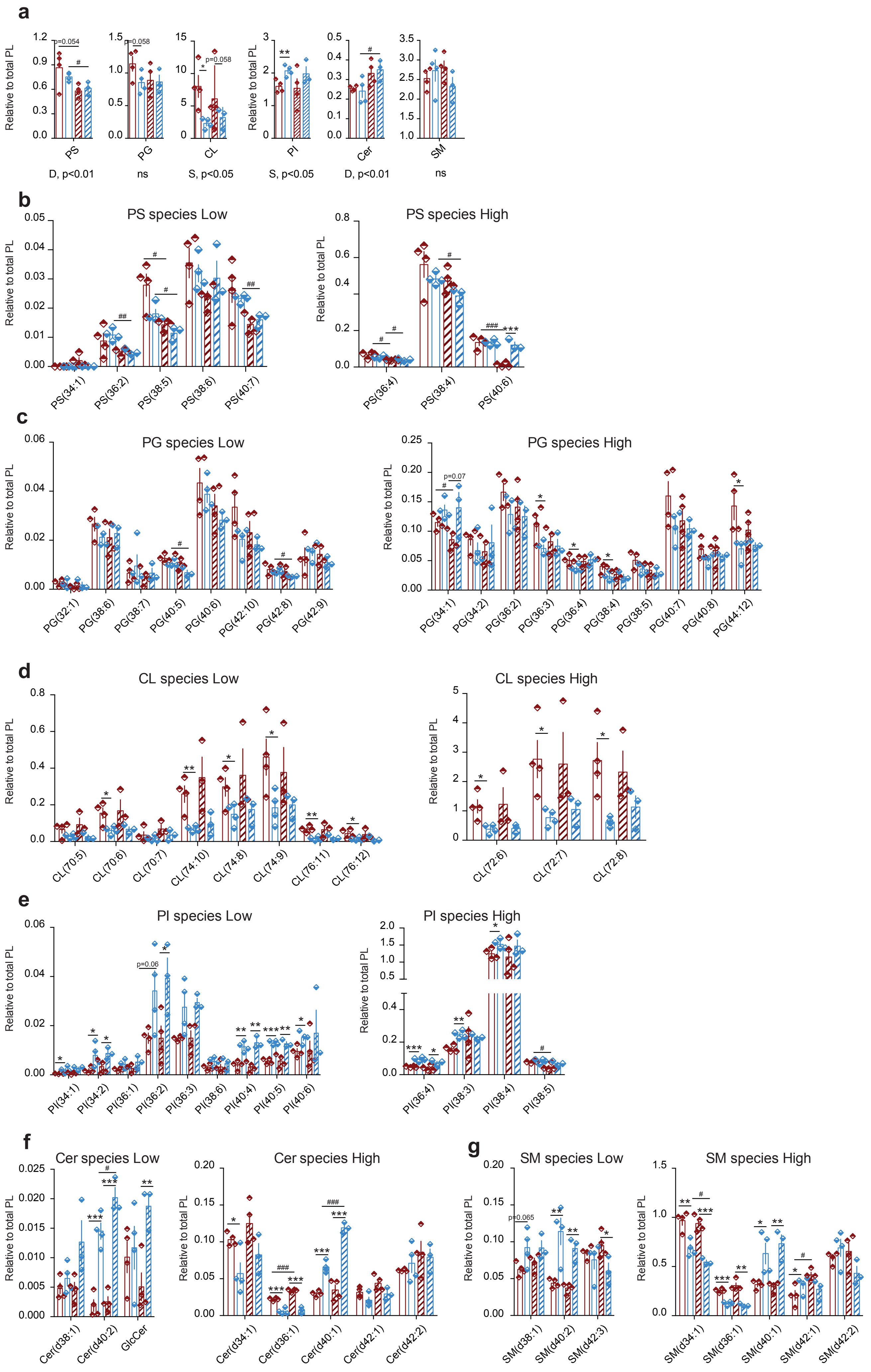


Supplementary Figure S5

a

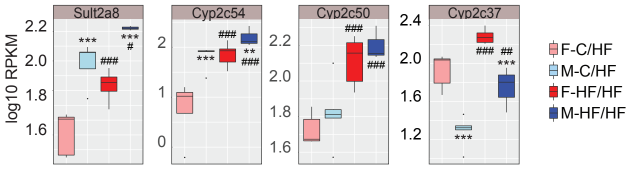


Supplementary Figure S6

Suppl.Table S1: Sex and MO effects on glucose, insulin and type II diabetes gene expression in offspring's liver.

| | SEX | | DIET | |
|---|---|---|---|---|
| **Pathway** | **C/HF** | **HF/HF** | **Females** | **Males** |
| Glycolysis / Gluconeogenesis | Acss2; Adh4; Ldha; Pklr; Aldh7a1; Dlat; Aldh3b3; Adh1; Akr1a1; Aldh1b1; Adh5; G6pc; Pck1 | Adh4; Pdhb; Aldob; Aldh1b1 | Acss2; Pklr; Gapdh; Dlat; Bpgm | Aldh1b1 |
| MAPK signaling pathway | Ntrk2; Egfr; Fgfr3; Mknk2; Tnfrsf1a; Rac2; Jund; Ddit3; Hspb1; Hspa8 | Rasgrp2; Ntrk2; Egfr; Nras; Mapk3; Tnfrsf1a; Il1r1; Hspb1 | Nf1; Ntrk2; Mknk2 | - |
| cAMP signaling pathway | Gabbr2; Rapgef4; Calm2; Rac2; Pik3r1; Creb3l3; Sox9; Acox1; Pde4b; Atp2a2 | Rapgef4; Gabbr2; Calm2; Mapk3; Acox1; Pde4b | Calm2; Pde3b | Fxyd1 |
| PI3K-Akt signaling pathway | Efna1; Egfr; Fgfr3; Prlr; Fn1; Vtn; Gng11; Sgk1; Hsp90aa1; Hsp90ab1; Pck1; G6pc; Ccnd1; Cdkn1a; Ccnd3; Creb3l3; Rxra; Pik3r1 | Egfr; Nras; Mapk3; Ghr; Prlr; Fn1; F2r; Prkaa2; Eif4e; Them4; Hsp90aa1; Gsk3b; Gys2; Ccnd1; Ywhaz | Efna1; Col4a1; Vtn; Ccnd1; Rxra | Hsp90ab1 |
| AMPK signaling pathway | Hmgcr; Fasn; Acaca; Ccnd1; Scd1; Lepr; Creb3l3; Pck1; G6pc; Srebf1; Stradb; Adipor2; Pik3r1; Acacb | Ccnd1; Gys2; Stradb; Prkaa2; Pparg | Acaca; Hmgcr; Scd1; Ccnd1; Adipoq; Fasn; Lep; Acacb | - |
| Insulin signaling pathways & Type II diabetes mellitus | Pik3r1; Ppp1r3b; Calm2; Pygl; Srebf1; Acaca; Acacb; Fasn; Pklr; G6pc; Pck1; Mknk2; Slc2a2/Glut2; Creb3l3; Rapgef4; Tnfrsf1a; Slc27a2/Fatp2; Pygl; Mlxipl/Chrebp | Gsk3b; Gys2; Calm2; Prkaa2; Eif4e; Nras; Mapk3; Rapgef4; Tnfrsf1a | Ppp1r3c; Calm2; Pde3b; Acaca; Acacb; Fasn; Pklr; Mknk2; Slc27a5/Fatp5; Adipoq | - |

DEG analysis using FDR<0.1 and p-value< 0.05 as a cut-off, that belong to the selected KEGG pathways in offspring's liver. For F-C/HF, n=5; M-C/HF, n=5; F-HF/HF, n=6 and M-HFHF, n=3.

Suppl.Table S2: Sex and MO effects on lipid metabolism gene expression in offspring's liver.

| | SEX | | DIET | |
|---|---|---|---|---|
| **Pathway** | **C/HF** | **HF/HF** | **Females** | **Males** |
| Biosynthesis of unsaturated fatty acids | Acot3; Elovl6; Scd1; Acot4; Acaa1b; Acox1; Acaa1a; Fads1; Acot1; Baat; Pecr | Acot3; Acaa1b; Hsd17b12; Acox1 | Elovl6; Scd1; Acot3; Fads1; Fads2 | - |
| Fatty acid biosynthesis | Fasn; Acaca; Acacb; Acsl5; Acsl3; Acsl1 | Acsl4 | Acaca; Acacb; Fasn; Acsl3; Acsl4; Acsl5 | - |
| Fatty acid degradation | Cyp4a12a; Adh4; Cyp4a12b; Acat2; Aldh7a1; Acsl5; Acaa1b; Acsl3; Echs1; Acox1; Acsl1; Cyp4a32; Cpt2; Acaa1a; Adh1; Acadm; Aldh1b1; Acadsb; Adh5 | Cyp4a12a; Cyp4a12b; Adh4; Acaa1b; Cyp4a32; Aldh1b1; Acox1; Acsl4 | Acat2; Acsl3; Acsl4; Acsl5 | Aldh1b1 |
| Fatty acid elongation | Elovl3; Acot3; Elovl6; Acot4; Echs1; Acot1 | Elovl3; Acot3; Hsd17b12 | Elovl6; Acot3 | - |
| Fat digestion and absorption | Acat2; Plpp3; Scarb1; Abcg8; Plpp1; Dgat2; Abcg5 | Apob | Acat2; Scarb1 | - |
| Non-alcoholic fatty liver disease (NAFLD) | Tnfrsf1a; Pik3r1; Rxra; Srebf1; Mlxipl; Pklr; Lepr; Adipor2; Xbp1; Ddit3; Casp7; Ndufa4; Cox4i1; Cox6a1 | Tnfrsf1a; Gsk3b; Prkaa2; Xbp1; Eif2s1; Cycs; mt-Co1 | Rxra; Pklr; Lep; Adipoq; Cyp2e1; Uqcr11; Cox8b; | - |
| PPAR signaling pathway | Fabp5; Cyp4a12a; Scp2; Pltp; Cyp4a12b; Scd1; Me1; Ubc; Acsl5; Acaa1b; Acsl3; Acox1; Acsl1; Cyp4a32; Cpt2; Acaa1a; Pck1; Plin5; Cyp8b1; Slc27a2/Fatp2; Acadm; Rxra; Angptl4 | Cyp4a12a; Cyp4a12b; Scp2; Acaa1b; Fabp5; Cyp4a32; Acox1; Me1; Ucp1; Pparg; Acsl4 | Me1; Fabp5; Scd1; Adipoq; Acsl5; Acsl3; Lpl; Pltp; Slc27a5/Fatp5; Rxra; Cyp7a1; Acsl4; Fabp4; Fads2 | - |
| Adipocytokine signaling pathway | Tnfrsf1a; Acsl1; Acsl3; Acsl5; Lepr; Pck1; G6pc; Rxra; Adipor2; Acacb | Tnfrsf1a; Acsl4; Prkaa2 | Acsl4; Acsl3; Acsl5; Lep; Rxra; Adipoq; Acacb | - |
| Regulation of lipolysis in adipocytes | Pik3r1; Plaat3 | | Fabp4; Pde3b | - |

DEG analysis using FDR<0.1 and p-value< 0.05 as a cut-off, that belong to the selected KEGG pathways in offspring's liver. For F-C/HF, n=5; M-C/HF, n=5; F-HF/HF, n=6 and M-HFHF, n=3.

Suppl.Table S3: Sex and MO effects on inflammatory gene expression in offspring's liver.

| | SEX | | DIET | |
|---|---|---|---|---|
| **Pathway** | **C/HF** | **HF/HF** | **Females** | **Males** |
| Apoptosis | Tnfrsf1a; Cflar; Casp6; Casp7; Dffa; Ddit3; Ctsd; Ctsh; Ctsl; Ctss; Pik3r1 | Tnfrsf1a; Casp6; Cycs; Eif2s1; Ctsb; Ctsc; Nras; Mapk3 | Ctsb; Ctsc; Ctsd | - |
| Natural killer cell mediated cytotoxicity | H2-T23; H2-K1; Rac2; Pik3r1 | Mapk3; Tyrobp; Fcer1g; Zap70; Nras | - | - |
| T cell receptor signaling pathway | Pik3r1 | Zap70; Mapk3; Nck1; Nras; Gsk3b | - | - |
| B cell receptor signaling pathway | Pik3r1; Rac2 | Mapk3; Gsk3b; Nras | - | - |
| TNF signaling pathway | Tnfrsf1a;Cflar; Creb3l3; Casp7; Il15; Pik3r1 | Tnfrsf1a; Mapk3; Cxcl1; Mmp14 | Ifi47 | - |
| Leukocyte trans endothelial migration | Pik3r1; Rapgef4; Rac2 | Rapgef4 | Cldn1 | - |
| Chemokine signaling pathway | Ccl9; Pik3r1; Rac2; Gng11; Elmo1 | Cxcl1; Cxcl9; Ccl9; Nras; Mapk3; Gsk3b; Rasgrp2 | Stat1 | - |
| NF-kappa B signaling pathway | Tnfrsf1a; Cflar | Zap70; Il1r1; Tnfrsf1a; Lbp | Lbp | - |

DEG analysis using FDR<0.1 and p-value< 0.05 as a cut-off, that belong to the selected KEGG pathways in offspring's liver. For F-C/HF, n=5; M-C/HF, n=5; F-HF/HF, n=6 and M-HFHF, n=3.

Suppl.Table S4: Sex and MO effects on hepatocellular carcinoma in offspring's liver.

| | SEX | | DIET | |
|---|---|---|---|---|
| **Pathway** | **C/HF** | **HF/HF** | **Females** | **Males** |
| Cell cycle | Ccnd1; Ccnd3; Hdac1; Cdkn2c; Cdkn1c; Cdkn1a | Ccnd1; Gsk3b; Ywhaz; Anapc11; Stag2 | Ccnd1 | - |
| Chemical Carcinogenesis | Cyp2c40; Gstp1; Cyp2c68; Cyp2c37; Cyp3a41a; Ephx1; Cyp3a44; Sult1a1; Cyp2c23; Cyp3a16; Cyp2c38; Gsta2; Cyp2c54; Gsta4; Cyp3a41b; Cyp2c39; Cyp2c70; Gstt2; Gsta1; Gstp2; Mgst1; Gstt3; Gstk1; Sult2a1; Sult2a3; Hsd11b1; Ugt2b5; Ugt1a2; Ugt1a6a; Ugt1a9; Ugt1a10; Ugt1a7c; Ugt1a5; Ugt2b1; Ugt1a6b; Ugt1a1; Ugt2b36; Cyp2b13; Cyp2b9; Aldh3b3; Adh1; Adh4; Adh5; Kyat3; Kyat1 | Cyp2c40; Gstp1; Cyp2c38; Sult1a1; Cyp3a44; Cyp3a41a; Ephx1; Cyp2c68; Cyp2c70; Cyp3a16; Cyp2c37; Cyp2c39; Gstt1; Mgst3; Mgst1; Gstm7; Gstt3; Sult2a1; Sult2a3; Hsd11b1; Ugt2b5; Ugt1a2; Ugt1a6a; Ugt1a9; Ugt1a10; Ugt1a7c; Ugt1a5; Ugt2b1; Ugt1a6b; Ugt1a1; Ugt2b34; Ugt2b36; Ugt2b38; Ugt2b35; Cyp2b13; Cyp2b9; Adh4 | Cyp2c23; Cyp2c50; Cyp2c37; Cyp2c54; Cyp1a2; Cyp2c38; Cyp2c29; Gsta4; Ugt1a2; Ugt1a9; Ugt1a10; Ugt1a7c; Ugt1a6b; Ugt1a1; Cyp2e1; Cyp2b9 | Cyp2c37; Cyp2c50; Cyp2c54; Ugt2b5; Ugt2b36 |
| Choline metabolism in cancer | Egfr; Pik3r1; Rac2; Chka; Chpt1; Plpp1; Plpp3 | Egfr; Mapk3; Nras; Chpt1 | | |
| MicroRNAs in cancer | Ccnd1; Egfr; Abcb1a; Cdkn1a; Fgfr3 | Egfr; Ccnd1; Pdcd4; Nras | Ccnd1; Slc45a3 | - |
| Mineral absorption | Slc11a2; Trf; Slc39a4; Steap2; Mt1 | Trf; Slc39a4 | Trf | - |
| mTOR signaling pathway | Pik3r1 | Eif4e; Mapk3; Prkaa2 | | - |
| Notch signaling pathway | Psenen; Hdac1 | Psen2; Hes1 | | |
| Oocyte meiosis | Ar; Calm2 | Cpeb2; Mapk3; Ywhaz; Anapc11; Calm2 | Calm2 | - |
| p53 signaling pathway | Ccnd1; Cdkn1a; Ccnd3 | Ccnd1; Cycs | Ccnd1 | |
| Pathways in cancer | Ccnd1; Wnt5b; Agtr1a; Gng11; Fn1; Pik3r1; Cdkn1a; Egfr; Fgfr3; Rac2; Rxra; Hdac1; Elob; Epas1; Ar; Hsp90aa1; Hsp90ab1 | Ccnd1; Gsk3b; F2r; Agtr1a; Fn1; Rasgrp2; Egfr; Nras; Mapk3; Dapk1; Pparg; Spi1; Cycs; Fh1; Hsp90aa1 | Ccnd1; Col4a1; Stat1; Rxra | Hsp90ab1 |
| Proteoglycans in cancer | Ccnd1; Esr1; Pik3r1; Egfr; Cdkn1a; Vtn; Fn1; Gpc1; Wnt5b; Ctsl | Ccnd1; Mapk3; Esr1; Nras; Ddx5; Egfr; Cav1; Cav2; Pdcd4; Sdc1; Fn1 | Ccnd1; Dcn; Cav1; Cav2; Sdc1; Vtn | |

| | | | | |
|---|---|---|---|---|
| Retinol metabolism | Cyp2b13; Cyp2c40; Cyp2a4; Cyp4a12a; Cyp2c68; Cyp2a5; Rdh11; Cyp2c37; Aox3; Ugt2b5; Cyp3a41a; Rdh16; Adh4; Cyp4a12b; Cyp3a44; Ugt2b1; Cyp2c23; Cyp3a16; Aox1; Cyp2c38; Ugt1a5; Cyp2b9; Cyp2c54; Cyp3a41b; Cyp4a32; Ugt1a2; Ugt1a10; Cyp2c39; Ugt2b36; Ugt1a6b; Hsd17b6; Lrat; Ugt1a6a; Adh1; Ugt1a7c; Ugt1a9; Dhrs4; Cyp2c70; Ugt1a1; Aldh1a1; Retsat; Dhrs9; Adh5 | Cyp2b9; Cyp2b13; Cyp2a4; Cyp2c40; Cyp4a12a; Aox3; Cyp2c38; Ugt2b5; Cyp3a44; Cyp4a12b; Adh4; Cyp3a41a; Cyp2c68; Ugt2b38; Cyp2c70; Cyp3a16; Ugt2b36; Aldh1a1; Ugt2b1; Aox1; Cyp2c37; Ugt2b35; Cyp2a5; Rdh9; Aldh1a7; Ugt1a5; Cyp2c39; Ugt1a1; Ugt1a2; Ugt1a10; Ugt1a9; Ugt1a6b; Ugt1a7c; Dhrs4; Ugt2b34; Dhrs3; Ugt1a6a; Cyp4a32 | Cyp2a4; Rdh16; Cyp2c23; Cyp2c50; Cyp2c37; Rdh9; Cyp2c54; Cyp1a2; Ugt1a1; Hsd17b6; Rdh11; Dhrs3; Cyp2b9; Aldh1a7; Cyp2c38; Cyp2a12; Ugt1a6b; Cyp2c29; Ugt1a7c; Ugt1a2; Ugt1a10; Ugt1a9 | Cyp2c37; Cyp2c50; Ugt2b5; Cyp2c54; Ugt2b36 |

DEG analysis using FDR<0.1 and p-value< 0.05 as a cut-off, that belong to the selected KEGG pathways in offspring's liver. For F-C/HF, n=5; M-C/HF, n=5; F-HF/HF, n=6 and M-HFHF, n=3.
